# Shielded by the dead: how killed bacteria shape the dynamics and evolution of innate immunity

**DOI:** 10.64898/2026.02.05.704043

**Authors:** Stephen P. Ellner, Nicolas Buchon, Tobias Dörr, Misha I. Kazi, Brian P. Lazzaro, Alexander Vladimirsky

**Affiliations:** Department of Ecology & Evolutionary Biology, Cornell University; Department of Entomology, Cornell University; Department of Microbiology, Cornell University; Cornell Institute for Host-Microbe Interactions and Disease; Weill Institute for Cell and Molecular Biology, Cornell University; Department of Mathematics, Cornell University

**Keywords:** host-pathogen interactions, innate immunity, immune system evolution, antimicrobial peptide

## Abstract

The outcome of an infection is determined by the dynamic interplay between microbial growth and host immunity. During a bacterial infection, bacteria killed by innate immune effectors can accumulate as corpses in the extracelluar space, where they can continue to bind (and thus sequester) immune effectors. The impacts on infection outcomes of continued biochemical activity (“sponginess”) by the dead have been generally overlooked in theoretical and empirical studies of within-host disease dynamics. We develop a mechanism-based mathematical model of within-host dynamics that incorporates host microbial sensing, the production of immune effectors, the interaction of those effectors with microbes, and shutdown of the immune response after an infection has been controlled. Corpse sponginess impedes the host’s ability to control infection, but at the same time, the rapid mopping up of effectors by bacterial corpses also protects host tissue against autoimmune self-harm from immune effectors still circulating after the infection has been resolved. This dual impact of bacterial sponginess alters the trade-off between damage done by infecting bacteria versus autoimmune damage, consequently shifting the evolutionarily optimal immune activation and shutdown kinetics. Thus, the sponginess of bacterial corpses likely shapes both short-term infection dynamics and the long-term evolution of immune systems.

## Introduction

The outcome of an infection is determined by the interplay of mutual negative feedbacks between host and pathogen. Innate immune interactions are biochemical in nature. For example, a primary defense against bacterial infection in barrier epithelia of animals, and of systemic defense in invertebrates, is production of antimicrobial peptides (AMPs) with direct antibiotic activity. A stereotypical mode of AMP action is to bind to pathogen cell membranes and aggregate to form open pores that, in sufficient numbers, result in osmotic leakage and cell death (Kleino and Silverman, 2014). Pathogenic bacteria have means of defending themselves against AMPs, including cell membrane alterations that reduce sensitivity to AMPs and secretion of proteases to degrade AMPs in the extracellular space (Joo et al., 2016).

An important but generally overlooked consequence of these direct biochemical interactions is that dead pathogens can still participate in the battle. This ongoing effect of dead cells may have important consequences for the dynamics and outcome of an active infection, especially once the dead are abundant. An infection that initially proliferates but is subsequently controlled must generate an appreciable number of dead pathogen cells that will persist in the host until they decompose completely or are filtered out, so there are likely to be times when killed pathogens outnumber the living. Dead pathogen cells or cell fragments may sustain the host immune response by releasing molecules that stimulate the response (Hixson et al., 2024; Troha et al., 2018). However, because the cell envelope is not immediately completely destroyed upon cell death, the corpse envelope (or envelope fragments) can also act as a “sponge” for AMP binding, diverting AMPs that otherwise could bind to living bacteria (Savini et al., 2020; Snoussi et al., 2018; Wu and Tan, 2019). In time-dependent killing experiments, we found that heat-killed bacteria protect live bacteria from AMP lethality in a dose-dependent manner (Fig. 1A). Killed cells can be substantially more attractive to AMPs than living cells (Savini et al., 2020; Snoussi et al., 2018), possibly because they no longer actively deploy anti-AMP defenses and because intracellular components may be accessible to AMP binding after the outer membrane is breached. Futhermore, once an infection is controlled, sequestration of circulating AMPs by the dead may reduce the autotoxic effects on the host of AMPs circulating at high concentration (Lei et al., 2019).

**Figure 1.**
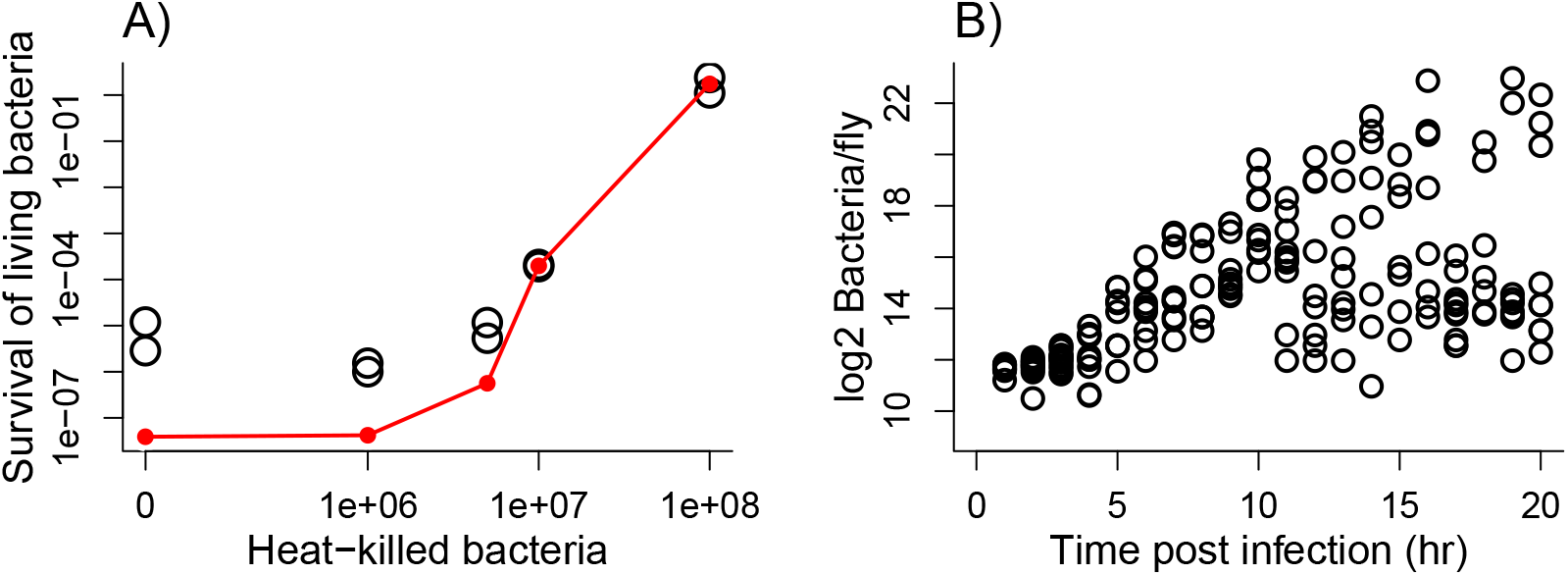
**A)** Presence of heat-killed bacteria increases the survival of living bacteria in the presence of AMPs. *Serratia marcescens* cells were exposed to the AMP cecropin B for 1 hour with or without the addition of heat-killed cells as indicated on the *x*-axis. Open black circles are experimental results (see below in section *Experimental demonstration of shielding by heat-killed bacteria*); closed red circles are model predictions at baseline parameter values. The nearly constant survival across the three lowest doses of heat-killed bacteria in the experiments is interpreted as a subpopulation of inert cells that neither divide nor die in the presence of AMPs. This subpopulation is too rare to affect the dynamics we consider (≈ 3 in a population of 2^18^ bacteria) and is not included in our model. **B)** Experimental infections of *Drosophila melanogaster* strain Canton S with the bacterium *Providencia rettgeri*. Each open circle represents total bacteria count for a fly sacrificed at the plotted time. Replotted using data from Fig. 3D of Duneau et al. (2017). The ≈ 2^10^ − 2^14^ (1000 - 10,000) surviving bacteria in the lower arm of infection outcomes at 10-20 hr post infection represent the empirically observed chronic infection (e.g., (Chambers et al., 2019; Duneau et al., 2017; Haine et al., 2008; Hidalgo et al., 2022; Jent et al., 2019; Kutzer and Armitage, 2016)). These cells are relatively inert and interact with the host weakly if at all (Ellner et al., 2021), and are omitted from this paper’s model (see main text). Figure made by script `Figure1_and_3.R.`

Our goal in this paper is to explore theoretically the impacts of the continued presence of killed pathogen cells on the progression and outcome of an infection. Based on the *in vitro* evidence that killed bacteria can absorb AMPs, we hypothesized that the dead cells generated in the course of fighting a pathogen could substantially alter the infection dynamics and outcome. We also explore some possible consequences of these impacts for evolution of the host immune response.

Our dynamic model describes a pathogenic bacterial population growing in an invertebrate host that uses AMPs as its primary immune defense, while the bacteria can respond by producing proteases that degrade AMPs. Parameter values for our numerical experiments are motivated by empirical studies with *Drosophila melanogaster* as the host (Duneau et al., 2017) and several opportunistic bacterial pathogens where the infection outcome is not a foregone conclusion: in some host individuals the bacteria overcome immune defenses and proliferate until the host dies, while other individuals eliminate the bacteria or reduce them to nearly harmless levels ((Duneau et al., 2017; Ellner et al., 2021; Frenkel et al., 2021), see Fig. 1B). We explore these possible impacts of killed bacteria on two time scales. During the course of a single infection, how do continued stimulation of the immune response and AMP binding by dead microbes affect the dynamics of infection and immunity in an individual host? And over the longer time scale of evolution, how might continued biochemical activity of dead bacterial cells drive adaptation of the host immune response, ensuring that the host can both control infection by living bacteria and deactivate AMP production effectively when it is no longer needed? The last two tasks are not well aligned, and lead to unavoidable trade-offs between minimizing pathogen damage and minimizing autotoxicity. We examine these trade-offs by considering the set of *“Pareto optimal”* immune responses, which will be described in detail below in **Materials and Methods**.

### Model Development

Our model aims to represent the host’s induced immune response and its consequences when the host faces a potentially lethal septic infection. The initial time *t* = 0 in our model corresponds to the moment when an infection has become established as systemic and consequently triggers the signaling cascade(s) that lead to a systemic induced immune response. The bacterial population is large enough by then that it can be modeled deterministically using differential equations.

To explore potential roles of dead pathogen cells in infection kinetics and the evolution of the immune response, the model includes: (i) gradual up-regulation of host immune responses, including AMP production, once a potentially harmful pathogen has been detected; (ii) binding of AMPs to living bacteria (causing cell mortality) and to dead bacteria (sequestration, or “sponging”); (iii) shutdown of the immune response once an infection has been controlled. Because all of these processes interact with effects of AMP absorption by dead bacteria, our model aims to represent the essential features of the mechanisms underlying each of these.

In particular, the timing of immune response — how long does it take for AMP concentrations to ramp up, and how long does it then take for the bacterial death rate to increase — is life-or-death for a host fighting an opportunistic infection (Duneau et al., 2017; Lafont et al., 2022). Our model is therefore most detailed with regard to the mechanisms determining these time lags, including multistep up-regulation of AMP production and gradual increase in bacterial mortality as additional bound AMPs are accumulated. Here, we provide a summary (Fig. 2, Box 1) and an explanation of the main features of the model. Additional details can be found in Supplement *§§* S1-S5.

**Figure 2.**
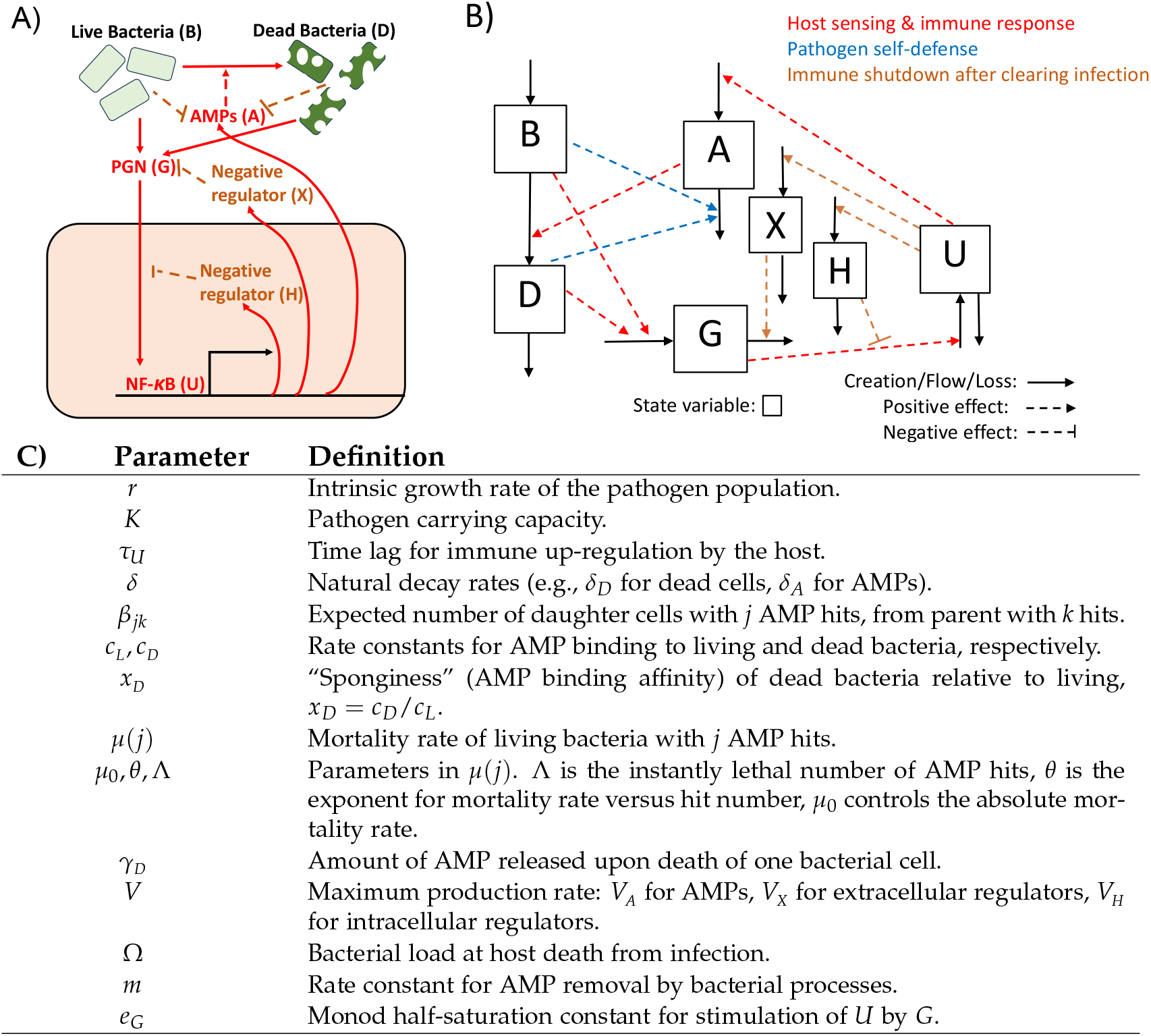
**A)** Diagram of the processes included in our model to predict the impacts of dead bacteria during an infection. **B)** Compartment diagram of the model. As described in the main text, the bacteria *B* are subdivided according to their number of AMP hits, and the up-regulation variable *U* is subdivided into *U*_1_, *U*_2_ and *U*_3_ roughly corresponding to transcription, translation, and secretion out of the host cell. Other state variables are dead bacteria *D*, AMPs *A*, peptidoglycan PGN *G* (bacterial products that stimulate the immune response), extracellular negative regulators *X* and intracellular negative regulators *H*. Solid arrows show material flows (creation, flows between compartments, loss), dashed arrows show effects of state variables on flow rates, color-coded according to their function. Note that a positive effect on a loss rate (e.g., effect of AMPs *A* on death rate of bacteria *B*) will have a negative effect on the affected state variable. **C)** Model parameters and their definitions. Parameter values are provided and justified in Supplement *§*S3 and *§*S5.

Fig. 2A shows the key processes included in our model. The state variables are living bacteria *B*, dead bacteria *D*, peptidoglycan (PGN) *G* produced by living and dead bacteria (Kleino and Silverman, 2014), AMPs *A*, host-derived extracellular negative regulators *X*, intracellular negative regulators *H*, and *U* representing the degree of host immune up-regulation driving production of *A, X*, and *H*. The difference between *X* and *H* is that *X* acts outside host cells, degrading the PGN that accumulates during a growing infection, while *H* acts within host cells, diminishing the degree of up-regulation in response to a given PGN concentration. In reality, hosts produce a mixture of different AMPs as well as different intra- and extra-cellular negative regulators but we simplify by combining these into three compartments (*A, X*, and *H*) rather than modeling multiple proteins that have qualitatively similar functions.

#### Box 1 Rescaled dynamic equations for the model

Time is scaled relative to bacterial doubling time in the absence of resource limitations (implying *r* = ln 2); rescalings of some state variables are described in Supplement *§*S1.4. *B*_*j*_ denotes the population of bacteria (cells per host) with *j* bound AMPs, *B* = *B*_0_ + … + *B*_Λ_ is the total population of bacteria. *W* represents the gradual decrease in immune response capabilities when bacterial load approaches Ω, the bacterial load upon death (BLUD). *W*(*x*) ≈ 1 for *x* ≈ 0, and *W* is monotonic decreasing with *W*(*x*) = 0 for *x* ≥ 1, to ensure that no AMPs or negative regulators are produced by dead hosts.

—— **Living bacteria cells with *j* AMP hits** (*j* = 0, 1, …, Λ) ——

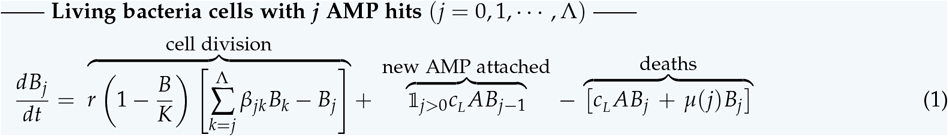

—— **Dead cells ——**

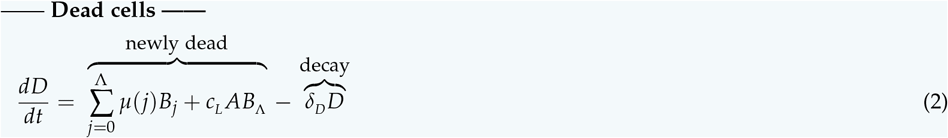

—— **Peptidoglycan, PGN ——**

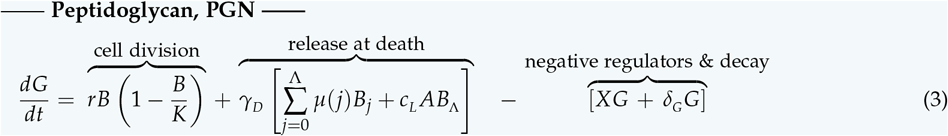

—— **Immune up-regulation ——**

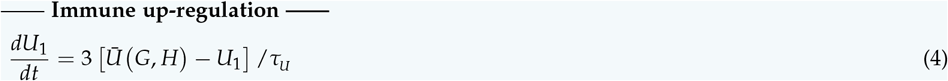

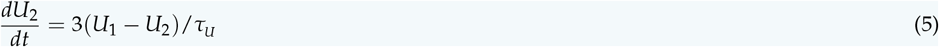

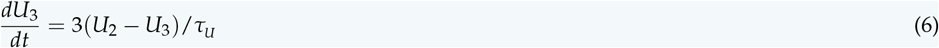

—— **Antimicrobial peptides, AMP ——**

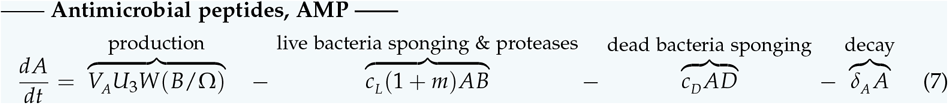

—— **Extracellular negative regulators ——**

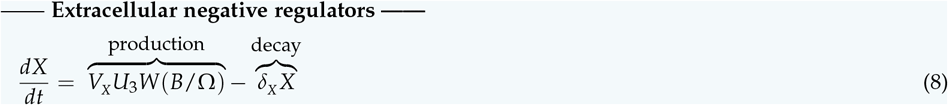

—— **Intracellular negative regulators ——**

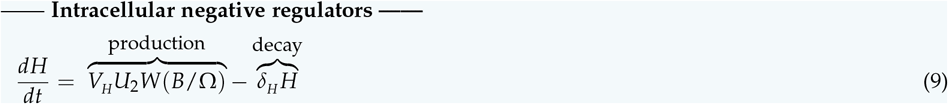

The system of ordinary differential equations (1)-(9) in Box 1 describes the full model of host-pathogen interactions. A detailed derivation of this model is given in Supplement *§*S1; here we outline the main assumptions and features of the model.

### AMP binding and lethality

We model AMP-induced pathogen mortality with a multi-hit model (Abel Zur Wiesch et al., 2015; Clarelli et al., 2020; Hemez et al., 2022; Teimouri et al., 2021) with bacteria classified by the number of AMP molecules bound to them. AMP-induced mortality occurs through creation of leaky pores in the cell envelope, which requires multiple AMP molecules acting in concert. In our model, the bacterial per-capita mortality rate *µ* increases monotonically with the number of AMPs bound to the cell, up to a threshold number Λ of bound AMPs where bacterial cell death is instantaneous. This results in a delayed and gradual increase of bacterial mortality following host immune up-regulation, which is realistic for physiologically reasonable AMP production rates, and magnifies the number of cells that must be killed to control an infection.

### AMP dilution

The number of AMPs bound to a live bacterium does not increase monotonically over time: in our model, when a cell divides, each AMP bound to it is equally likely to end up in either of the two daughter cells, independent of the fate of other bound AMPs. For each division of a cell with *k* AMP hits, the expected number of daughter cells with *j* ≤ *k* hits is given by eqn. (S2) in Supplement *§*S1.2.

### Bacteria death

Because bacteria cell death results from lysis, we assume that the PGN from a killed cell becomes immediately accessible for recognition by the host at the moment of cell death. The remaining bacterial corpses (or corpse fragments) continue to sponge away AMPs for a much longer period. The “sponginess coefficient” *x*_*D*_ = *c*_*D*_ /*c*_*L*_ (i.e., the relative rate of AMP attachment to dead versus live bacteria) is a main focus of our attention throughout this paper.

### Bacteria defense

Some AMPs are also degraded by proteases that are produced by the bacteria. Following Ellner et al. (2021) we make the quasi-steady-state assumption that protease concentration is proportional to bacterial abundance. We assume that for each AMP that binds to a live bacterium *m* others are destroyed by proteases.

### Host immune up-regulation

Thepeed and intensity of host immune response depend on the intensity of the stimulus. The immune up-regulation *U* is modeled as a multi-step process (using state variables *U*_1_, *U*_2_, and *U*_3_, corresponding roughly to transcription, translation, and secretion out of the host cell) in order to capture the time delay *τ*_*U*_ between signal (PGN) and response (AMP production). A detailed model of the Imd signaling cascade (see Supplement *§*S2) suggests that the steady-state relationship between PGN concentration and immune up-regulation (i.e., the amount of bound promoter Relish in the cell nucleus), with all else held constant, is approximately a Monod curve. We use *Ū* (*G, H*) = *Ge*^−*H*^ /(*e*_*G*_ + *Ge*^−*H*^) to represent the quasi-equilibrium level for *U*_1_ given the current concentration of PGN (*G*). While this quasi-equilibrium response grows with *G*, it decreases when there are more intracellular negative regulators *H*.

### Negative regulator production

Because *H* acts within the host cell, its production is driven by the current level of *U*_2_, while the production of AMPs and extracellular negative regulators *X* (responsible for removing PGN) are driven by *U*_3_. Thus, an increase in *H* indirectly suppresses the production of new AMPs as well as the production of new negative regulators of both kinds.

### Host death

All immune response activities also gradually shut down if the pathogen population approaches the lethal pathogen burden Ω, the bacterial load upon death (called BLUD). However, for modeling bacterial growth dynamics, we assume that the carrying capacity *K »* Ω is much greater because the bacteria can continue to multiply in a dead host.

### Natural decay

Dead bacteria, AMPs, PGN, and negative regulators *H* and *X* are all subject to natural decay at specified rates *δ*_*D*_, *δ*_*A*_, *δ*_*G*_, *δ*_*H*_, and *δ*_*X*_, respectively.

## Materials and Methods

### Experimental demonstration of shielding by heat-killed bacteria

In 1 mL of LB broth growth medium, 7 *×* 10^6^ live bacteria *Serratia marcescens* were introduced, followed immediately by addition of heat-killed bacteria and AMP cecropin at concentration of 20*µ*g per mL, approximately 10^15^ AMP molecules. Survival (number of living bacteria) was assessed after one hr. There were two replicates each, at five different initial doses of heat-killed bacteria: zero, 10^6^, 5 *×* 10^6^, 10^7^, and 10^8^. Survival of living bacteria was very low at the three lowest doses (under 0.002% survival). With 10^7^ heat-killed bacteria the survival of living bacteria was 0.02%, and with 10^8^ the number of living bacteria increased to 111% and 240% of initial number in the two replicates.

### General computational Methods

All numerical calculations were done in R version 4.2.0 or higher for Windows (R Core Team, 2025), using BLAS from the Intel oneAPI Math Kernel Library, version April 2023 or later.

Model solution trajectories were calculated numerically with the ode function in the **deSolve** R package (Soetaert et al., 2010). We generally used the default integration method lsoda, an adaptive solver that switches between stiff and non-stiff solvers based on an internal monitoring of system stiffness. For model calibration, which was very time-demanding, we used method vode with method flag `mf=22`. This specifies a stiff solver using backwards differentiation and finite difference approximation of the full Jacobian (Brown et al., 1989). For our model `vode` was generally faster, but more likely to fail (usually in step size adjustment) when unrealistic parameter values produced extreme model dynamics, generating an error message and rejection of those candidate values.

### Choice of default parameter values

We used two approaches to select the default parameter values for numerical studies of our model. The general approach is outlined here; full details are in Supplement *§*S5 and *§*S3.

The available information in the literature, or from our own experiments, is sufficient to identify a range of plausible values for many model parameters. For those, we used the *geometric mean principle* to select a single default value from the plausible range. Namely, when the plausible range for a parameter *θ* was *a* ≤ *θ* ≤ *b*, we selected 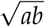 as its default value. This choice minimizes the maximum possible fold error (i.e., the worst-case ratio) between the chosen value and any other value in the plausible range. Baseline values were chosen this way for Ω, *K, τ*_*U*_, *γ*_*D*_, *m, x*_*D*_, *δ*_*D*_, *δ*_*A*_, *δ*_*X*_, *δ*_*H*_, *δ*_*G*_, and Λ. See Supplement *§*S3 for details.

The parameters *V*_*A*_, *V*_*X*_, *V*_*H*_, *c*_*L*_ and the parameters in the pathogen mortality function 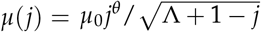 were estimated by model calibration, specifically they were chosen to minimize an error function combining the discrepancies between model simulations and three performance targets, when using the baseline values for the other parameters.

The first target was the bacterial survival rate in the *in vitro* experiments with shielding by 10^7^ and 10^8^ heat-killed bacteria (see below in *Experimental demonstration of shielding by heat-killed bacteria*). As noted in the Fig. 1 caption, the survival rates in the other treatments are interpreted as a very small number of inert bacteria (one in 10^5^–10^6^). These are omitted from our model because they would have no effect on the dynamics in the situations that we consider.

The second target was a pair of stylized trajectories of total living bacteria population dynamics (Fig. S7) representing typical behavior in the experiments in Duneau et al. (2017). The error function term from this target was the sum over the two trajectories of the mean absolute difference between the natural logs of observed and predicted total living bacteria.

The third target was the property that AMP concentrations decrease monotonically after reaching their maximum during the course of a controlled infection. This property has not been demonstrated experimentally. We nonetheless impose it here to rule out the very small window of parameter values very near the split between clearing the infection and succumbing to it, where model solutions oscillate at intermediate values for several days, a property that was either very rare or entirely absent in experiments (Duneau et al., 2017). The error function term from this target was the fraction of time, after AMP concentration reached its maximum, that *dA*/*dt* was positive between *t* = 0 and 72 hours.

Supplement *§*S5 provides details on how the error terms were computed and combined into an overall objective function, and how that function was optimized.

### Parameter sensitivity analysis

We evaluated the relative sensitivity of infection outcomes to changes in different model parameters by computing the parameter elasticities of four measures of the host’s ability to fight off infection and the costs of doing so. Our first measure, primarily highlighting the immune response effectiveness, is the maximum initial pathogen load (at *t* = 0, when the infection becomes systemic) that is controlled by the host, leading to host survival. The remaining three measures highlight different costs associated with controlling the pathogen. The second is the cumulative AMP production over time, given by 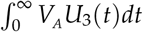 *dt* where *𝒰*_3_ measures the fractional up-regulation of immune effector secretion out of host cells, and *V*_*A*_ is the maximum possible production rate. The third is the cumulative auto-toxic damage to the host by AMPs, assumed proportional to 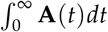 where **A** is the number of circulating AMP molecules (not including those bound to living or dead bacterial cells). Finally, the fourth measure is the cumulative direct damage to the host by the pathogen, assumed proportional to 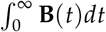 where **B** is the total number of living bacteria at time *t*.

Maximum initial load was approximated by finding the initial pathogen load such that the total number of bacteria after 48 hours was equal to the initial number. For the second and third measures, the infinite integrals were evaluated by integrating up to a finite time *T* such that *U*_3_ and *A* had fallen to zero and remained there for some time, because the host either had cleared the infection or was dead, and in either case all immune activity has ceased. Integrals were evaluated by computing model solution trajectories at a fine grid of time points (every 0.05 hr) and approximating the integrals by trapezoid rule.

The last two measures have the un-biological feature that if the pathogen kills the host, the host continues to accrue damage even after it is dead. However, post-mortem autotoxic damage is actually near zero, because a dead host produces no new AMPs and circulating AMPs are almost instantly absorbed by the rapidly growing pathogen population. Allowing damage from bacteria to accrue on the host has the desirable feature that being killed by infection is much more costly than being nearly killed but eventually controlling the infection.

For any response *Y* as a function of some parameter *θ* with baseline value *θ*_0_, the parameter elasticity of *Y* to *θ* is defined as the fractional change in *Y* resulting from a small fractional change in *θ* away from *θ*_0_, with all other model parameters held constant. We calculated elasticities using *±*2% changes in each parameter. The parameter elasticity is thus

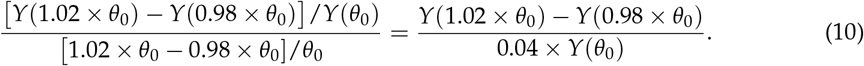

This formula was evaluated at our best-fitting parameter values, with initial pathogen dose of 3000 cells. That initial dose is close to the maximum controlled dose (≈ 3500 cells) but far enough away from the “split” between controlled and uncontrolled infections that 2% changes in model parameters did not produce uncontrolled infections.

In addition, we calculated how each response variable changed as a function of *x*_*D*_, the AMP affinity of dead relative to living bacteria, and maximum AMP production rate *V*_*A*_, over a fine grid of values from 50% to 150% of the default values of those parameters. This grid crosses the “split” in infection outcome, where some hosts launch on a trajectory toward death and others head toward survival. We found that as the split is crossed, the changes in the responses (all but the first above) were not monotonic at fine scales because model solutions very close to the split are oscillatory, and small parameter changes can make a big difference in when the pathogen population crashes to extinction, versus when it kills the host and commences uncontrolled growth.

### Pareto Front computations: “optimal” trade-offs in host immune response

In our evolutionary analysis, host performance is constrained by two mechanistically distinct components of harm generated by the within-host dynamics: cumulative pathogen damage (i.e., the time-integrated burden of living bacteria) and cumulative immune self-damage (i.e., the time-integrated toxicity of circulating AMPs). For any candidate immune strategy, here represented by evolvable parameters such as the maximal AMP production rate and intracellular negative regulation (i.e., *V*_*A*_ and *V*_*H*_), the model produces a corresponding pair (*𝒟*_*B*_, *𝒟D*_*A*_) of these two damages. Because increasing immune aggressiveness typically reduces bacterial damage while increasing AMP-toxicity damage, there is generally no single strategy that simultaneously minimizes both objectives. The Pareto front is therefore the set of non-dominated strategies, meaning you cannot “beat” any of them on both axes at once: for anything Pareto-optimal, there is no feasible strategy that yields lower bacterial harm and lower immune self-harm simultaneously. Equivalently, each point on the front represents a strategy such that any attempt to further decrease one type of damage necessarily increases the other. This construct is useful here because the true mapping from (𝒟_*B*_, 𝒟_*A*_) to fitness is rarely known *a priori* (i.e., how strongly selection penalizes bacterial harm versus immunopathology), and the Pareto front includes all those strategies that could be optimal for at least one plausible way of weighting these damages into overall fitness. In practice, we approximate portions of the Pareto front by repeating the optimization while varying how strongly selection penalizes bacterial harm versus immune self-harm, and compiling the resulting optimal strategies. Mathematically, this is essentially a *linear scalarization* method; see Supplement *§*S7 for full details. Our main conclusions hinge on how corpse sponginess shifts the achievable trade-off boundary (Fig. 5), rather than on any one particular choice of weighting.

## Results

### Infection time scale

The negative feedbacks between host and pathogen in our model allow either to get the upper hand – the host suppressing the infection in the early stages of proliferation, or the pathogen overwhelming the immune defense and killing the host. Consequently, our model exhibits the same bimodality of outcomes as previous models (in Ellner et al., 2021; Lafont et al., 2022; van Leeuwen et al., 2019). In particular, the model replicates the empirical observation in Duneau et al. (2017) for the *Drosophila* model system showing that variation among individuals in the time required for immune up-regulation can determine whether the host defeats an infection or succumbs to it (Fig. 3A). A greater delay in immune up-regulation causes a delay in AMP production (Fig. 3B). That delay allows the bacterial population to continue growing longer, reaching a tipping point where the host’s AMP production is outpaced by bacterial cell divisions producing new cells with fewer AMP hits each than the parent.

**Figure 3.**
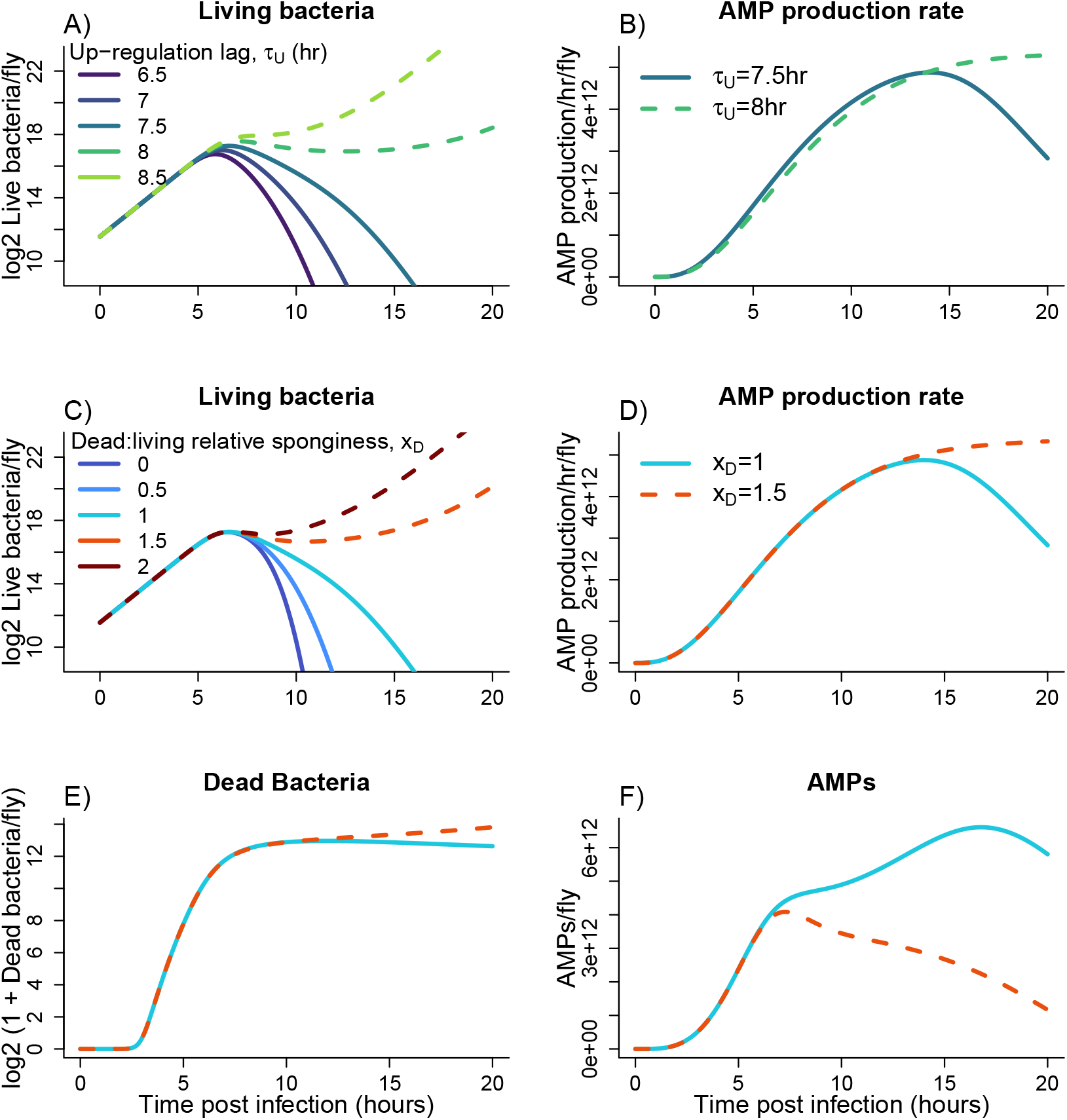
Immune up-regulation lag and AMP sponging by the dead can both be life-or-death for the host, but the mechanisms are completely different. In all panels, solid curves are scenarios where the pathogen is cleared, dashed curves are scenarios where the host is eventually killed. **A)** Model simulations capture the observed divergent infection outcomes (Fig. 1B) as the time-constant for immune upregulation, *τ*_*U*_, is varied with other parameters at their baseline values (baseline *τ*_*U*_ = 7.5 hrs). **B)** Increasing *τ*_*U*_ leads to a delay in AMP production, which allows bacterial population growth to continue longer and pass a tipping point. **C)** Model simulations also predict divergent outcomes when we vary *x*_*D*_, the AMP “sponginess” of dead cells relative to that of the living. **D)**,**E)**,**F)** How increased AMP sponging by the dead produces divergent outcomes. Even though AMP production rate is the same up to *t* ≈ 12, the accumulation of dead bacteria between *t* = 5 and *t* = 10 causes the AMP concentration to decrease starting at *t* ≈ 7 if the dead absorb AMPs faster, allowing bacterial population growth to resume and eventually kill the host. Figure made by `script_Figure1_and_3.R`.

We note that our model predictions depart from experimental data in that a defeated pathogen in the model is completely eliminated, whereas experimentally there is often a small population that survives the host immune response and persists as a low-level chronic infection having little or no effect on the host. It would be straightforward to make our model exhibit this feature by including a subpopulation of near-inert persister cells (the approach we took in Ellner et al. (2021)). However, doing so would not change our conclusions about the role of the dead bacteria, so we omit it here for the sake of simplicity.

Our model also makes the new prediction that the affinity of dead cells for binding by AMPs (their AMP “sponginess” relative to that of living cells, *x*_*D*_) can be a matter of life or death for the host (Fig. 3C). If the delay in immune upregulation is kept the same but we instead vary AMP sponging by the dead, an infection that would quickly be controlled if AMPs did not bind to killed bacterial cells (lowest curve in Fig. 3C) is controlled much more slowly if living and dead bacterial cells have equal affinity for AMPs (*x*_*D*_ = 1), and instead kills the host if dead cells have slightly higher affinity than living cells (*x*_*D*_ ≥ 1.5). Higher AMP affinity of dead cells has indeed been observed for *E. coli* interacting with human AMP LL-37 (Snoussi et al., 2018) and porcine AMP PMAP-23 (Savini et al., 2020). Converting one kind of adversary into another – i.e., turning a living bacterium capable of proliferating into a corpse that more quickly absorbs AMPs – may thus be a kind of Pyrrhic victory, by no means optimal from the host perspective. But hosts cannot avoid the constraint (Gould and Lewontin, 1979) that a killing creates a corpse.

Although very similar pathogen dynamics are produced by variation in immune up-regulation lag *τ*_*U*_ (Fig. 3A) and variation in dead relative sponginess *x*_*D*_ (Fig. 3C) the underlying mechanisms are very different. Increasing the relative sponginess from *x*_*D*_ = 1 (equal affinity) to *x*_*D*_ = 1.5 has near-zero effect on AMP production (Fig. 3D), or on the number of dead bacteria (Fig. 3E), up to *t* ≈ 12. But the resulting AMP concentrations diverge much earlier, around *t* = 6 (Fig. 3F), due to higher AMP absorption by the large number of dead bacteria.

This potential effect of dead bacteria on the build-up of AMP concentration, and how that can affect infection clearance, are the new features introduced in this paper. The capacity for this to occur in the model is a consequence of the experimentally demonstrated mechanisms that we included in the model, but whether or not it actually occurs depends on the combination of parameter values. For example, if the PGN released by the dead stimulated a higher host immune response, this could compensate for AMP sponging and give the host the upper hand. So it is significant that we find these effects at parameter values that were either based on estimates in the literature or calibrated to match the available data, rather than values that were chosen to achieve some particular consequences of differing levels of AMP binding by the dead.

Many model parameters contribute to the speed or strength of the host immune response and therefore contribute to determining whether the host controls or succumbs to the infection. Model sensitivity to small changes in parameter values (Fig. 4) confirms that time is of the essence for successful immune response (Duneau et al., 2017; Ellner et al., 2021). The response variables in Fig. 4 are four measures of the host’s ability to fight off infection and the costs of doing so; see *Methods: Parameter sensitivity analysis* for mathematical definitions of the response measures and their elasticities, and calculation methods. Bar height measures how strongly each parameter affects the response variable (named in each panel’s label), and whether the effect is positive or negative. For example in panel A, a 1% increase in *τ*_*U*_ would cause a 2.73% decrease in the maximum controlled initial dose. The model parameters omitted from the Figure had elasticity ≤ 0.2 for all response measures. Because elasticities are proportional changes, they are independent of the units in which the response is measured.

**Figure 4.**
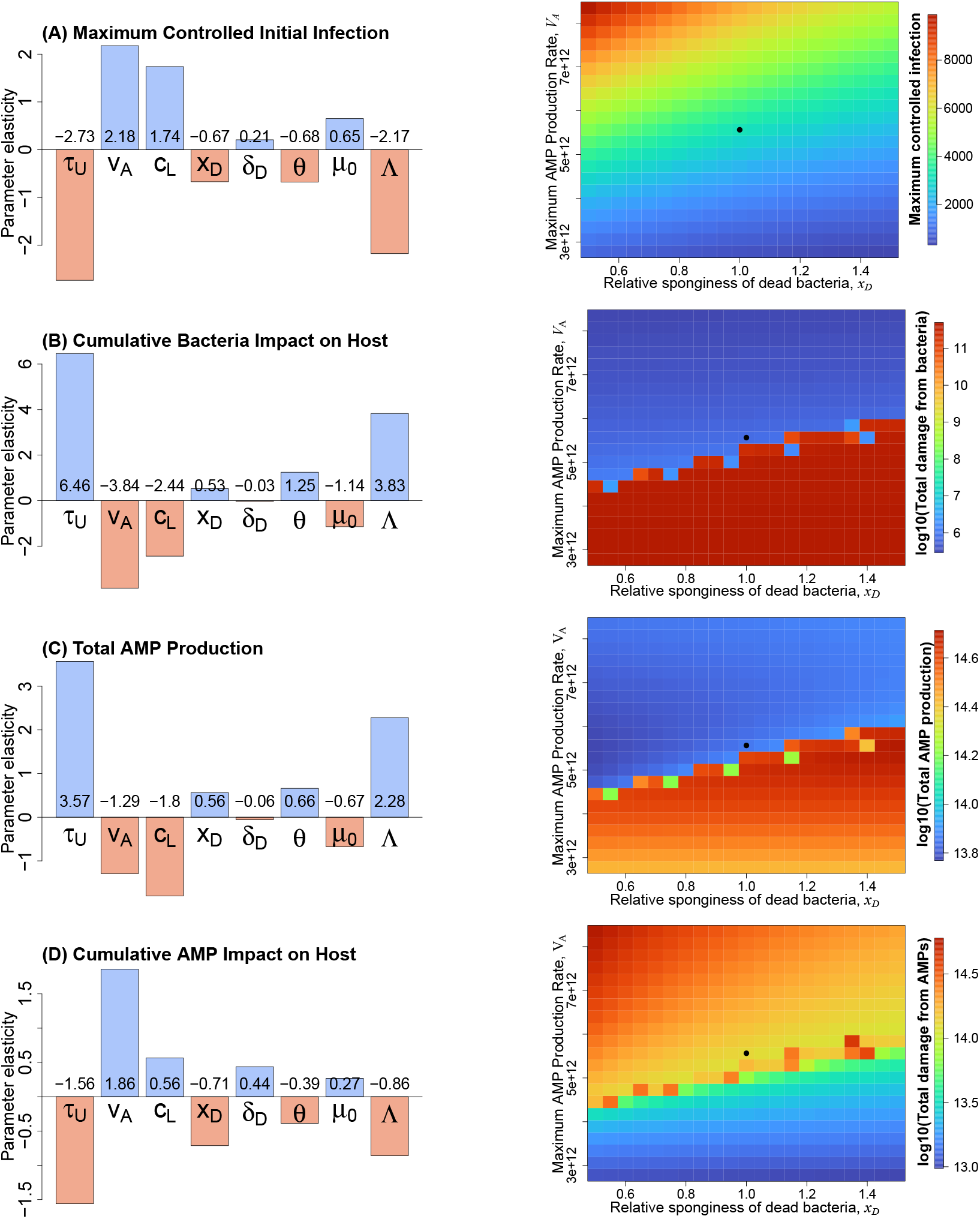
**Left column:** Parameter sensitivities for the model parameters (8 out of 18 total parameters) that had the largest impacts on four measures of host ability to control an infection or impacts on the host of a controlled infection. Plotted values are elasticities, the percent change in the response measure relative to small percent changes in parameter values (evaluated numerically by making *±* 2% parameter changes). For our baseline parameter values, the maximum controlled initial pathogen load is ≈ 3500 cells; panels (B)-(D) used initial load of 3000 cells. **Right column:** heat maps displaying how the response measures vary across a wider range of values for two model parameters, the relative AMP affinity of dead bacteria, *x*_*D*_ and the maximum AMP production rate by the host, *V*_*A*_ . Black solid circles indicate defaul parameter values. Figures made by R scripts in the Figure 4 code archive folder.

Slower immune up-regulation (large *τ*_*U*_) decreases the maximum controllable initial pathogen dose (panel A) and increases the cumulative damage done by the pathogen as well as the total AMP production (panels B and C). This is intuitive, as slower up-regulation of immune response allows a longer initial window for uncontrolled pathogen proliferation, resulting in more bacteria that do more damage and require more AMPs to control. Less intuitively, slower upregulation actually decreases the cumulative self-damage from AMPs (panel D) because higher AMP production is more than counter-balanced by the greater absorption of AMPs by the larger population of bacterial cells, both living and the corpses remaining after the infection has been controlled, thereby limiting autoimmune damage. Increasing bacterial tolerance of AMPs by increasing the instantaneously lethal dose Λ shows the same pattern as *τ*_*U*_ . As values of Λ increase, more time is required to control the pathogen. The maximum production rate of AMPs, *V*_*A*_, and the AMP binding affinity of living cells *c*_*L*_ show exactly the opposite pattern, because they hasten rather than delay the effective impact of immune up-regulation on the bacterial population.

The relative sponginess of dead cells, *x*_*D*_, also has substantial effects, following the same pattern as *τ*_*U*_ and Λ. By absorbing AMPs that could otherwise attack living pathogen cells, the dead act as a partial shield to the living, increasing the time (and the cumulative AMP production) needed to reverse pathogen proliferation. Yet at the same time, sponging of AMPs partially shields the host against the auto-toxic effects of circulating AMPs, decreasing cumulative AMP impact on the host. The pathogen corpses thus act as a double-edged sword for the host: infections are harder to control (i.e., faster immune induction or higher maximum AMP production rate is needed to compensate for AMP sponging by the dead), but the self-inflicted collateral damage from the AMPs required to control an infection is reduced. Thus the dead do not just shield their living relatives, they also shield the host.

### Evolutionary time scale

It is natural to expect hosts to co-evolve with narrowly specialized bacteria they are often exposed to. But with opportunistic (and broadly distributed) bacterial pathogens, it is a reasonable simplification to consider bacterial properties as fixed and to ask how the host’s immune mechanisms might evolve over time in response to frequent exposures. The assumption is that maximizing host fitness would correspond to minimizing the negative impacts of various costs associated with fighting off that infection. However, the mix of positive and negative impacts of dead bacteria on the host results in non-trivial and subtle implications for (a) which types of immune response deserve to be called “optimal” and (b) how these “optimal” responses depend on bacterial properties. By shielding living pathogens from AMP damage, the dead increase the importance for the host of having a high potential rate of AMP production. By removing many of the AMPs once the infection is controlled, the dead bacteria also shield the host from autotoxicity damage, reducing one cost of higher AMP production.

As an initial exploration of the evolutionary implications of shielding by the dead, we used our model to ask how the sponginess of the dead relative to the living affects the adaptively optimal values of two key parameters of the host immune response, *V*_*A*_ and *V*_*H*_ . (Of course, other parameter combinations could be similarly considered; see Supplement *§*S8 for an example.) Our purpose here is to test provisionally whether sponging of AMPs by bacterial corpses *might* have different (indeed, opposite) effects on the infection and evolutionary time scales.

*V*_*A*_ is the main determinant of the strength of the immune response once it is up-regulated, and *V*_*H*_ determines the speed and intensity of the first-acting component of the immune shutdown after the host clears an infection (extracellular regulators *X* come into play more slowly because they have to be secreted out of the cells in which they are produced, and they act more gradually because they gradually degrade PGN to reduce subsequent immune stimulation whereas *H* directly blocks the response to PGN).

We asked how these parameters could be expected to evolve in response to natural selection for minimizing two costs of infection: the cumulative damage to the host from bacteria (Fig. 4B) and the cumulative damage to the host from circulating AMPs (Fig. 4C). These are conflicting goals. If damage caused directly by bacteria was the only concern, infinite *V*_*A*_ and zero *V*_*H*_ would be optimal (instantaneous and infinite immune system activity). However, such a strategy would result in unsustainably high costs of AMP production and autoimmune damage (indeed, this is a leading explanation for why immune responses are inducible by stimulus instead of being constitutively active at high levels (Lazzaro and Tate, 2022)).

To explore evolution of immune system activity in response to these two kinds of damage, we computed the set of *Pareto optimal* host defense strategies for different values of *x*_*D*_, the sponginess of dead relative to living bacterial cells. A strategy is Pareto optimal if neither of these two types of damage can be further reduced without increasing the other (Miettinen, 1999). Such strategies provide us with a lower bound for the damages actually incurred by *any* evolved immune response mechanisms.

Pareto fronts, i.e., curves showing the incurred cost pairs (bacterial damage, AMP damage) for all Pareto-optimal immune response strategies, are shown in Fig. 5. Suppose that a host is willing to tolerate damage from bacteria up to some level, but no more than that. A vertical line that intersects the Pareto front at that value of bacterial damage indicates the lowest AMP damage that such a host can achieve while limiting bacterial damage to the level it is willing to accept. The dotted vertical line in Fig. 5 is an example.

**Figure 5.**
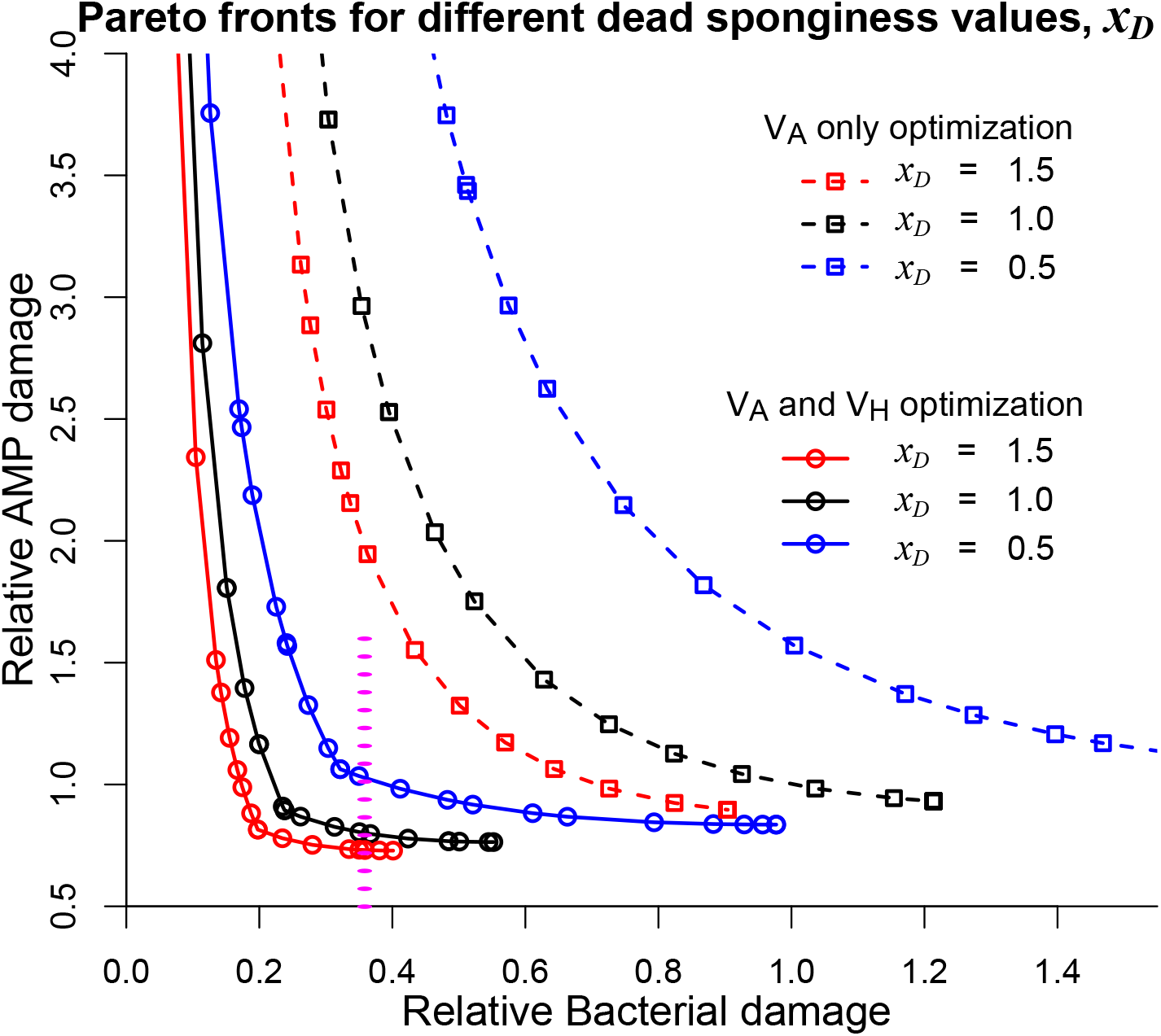
Sponginess affects the trade-off between damage from bacteria and damage from circulating AMPs, and shapes how the immune system should evolve to balance these costs. The host immune response has to balance direct damage caused by infecting bacteria against the collateral auto-toxic effects of circulating AMPs, including AMPs that persist after an infection is controlled. The displayed Pareto front (PF) curves show the non-dominated (Pareto optimal) immune responses such that neither damage can be further decreased without increasing the other. All damages are “relative”; i.e., shown as fractions of damage incurred during a controlled infection in the model with our baseline parameter values. Dashed curves correspond to optimizing the rate of AMP production (*V*_*A*_) only; solid curves correspond to optimizing the production rates of both AMPs and intracellular negative regulators (*V*_*A*_ and *V*_*H*_). Colors corresponds to dead bacteria “sponginess” relative to living bacteria (*x*_*D*_). The plotted Pareto-optimal strategies all lead to controlled infections; the full Pareto fronts also include biologically irrelevant strategies (outside the plotted region, and disconnected from the plotted curves) where there is no immune response and very high bacterial damage. The vertical dotted line indicates the level of bacterial damage studied in Fig. 6. Figure made by scripts in the Figure 5 code archive folder.

Such frontiers are useful for visually interpreting the shifting trade-offs in different Pareto-optimal immune responses. Moving to the left along the Pareto front from the dotted line intersection, it is easy to see how much of an increase in AMP damage will necessarily result if the host decreases the bacterial damage that it is willing to tolerate by any specific amount. The damages resulting from any evolved immune response would necessarily lie “northeast” of the corresponding Pareto front curve.

Whether we limit the host to optimizing the value of AMP production rate *V*_*A*_ (dashed curves) or also allow optimization of *V*_*H*_ (solid curves), Pareto optimal strategies for higher sponginess of dead bacterial cells relative to living ones (*x*_*D*_ = *c*_*D*_ /*c*_*L*_) are “southwest” of those for lower sponginess. This reflects that after an infection has been controlled, dead pathogens shield the host from subsequent autoimmune damage. Moreover, the ability to optimize both the speed of both activation and shut-down allows for an equally effective immune response with much lower resulting damages to the host (i.e., the solid lines are southwest of the dashed lines in Fig. 5).

The relative positions of the Pareto fronts for different levels of sponginess of dead bacterial cells relative to living ones shows that higher sponginess of the dead allows the host to achieve a lower AMP damage through optimization of its immune response for any given level of damage from bacteria. In addition, the tolerated bacterial damage is achieved through different immune response strategies, producing a different time-course of infections (Fig. 6). For example, for the lowest AMP sponginess (blue Pareto front), PGN lingers much longer and would stimulate continued immune up-regulation, except that this is blocked by much higher level of intracellular regulators *H* that prevent an unnecessary and damaging continuation of AMP production. AMPs nevertheless also linger longer because fewer are removed by dead bacteria, so the overall damage from AMP toxicity is higher than when the dead are more spongy.

**Figure 6.**
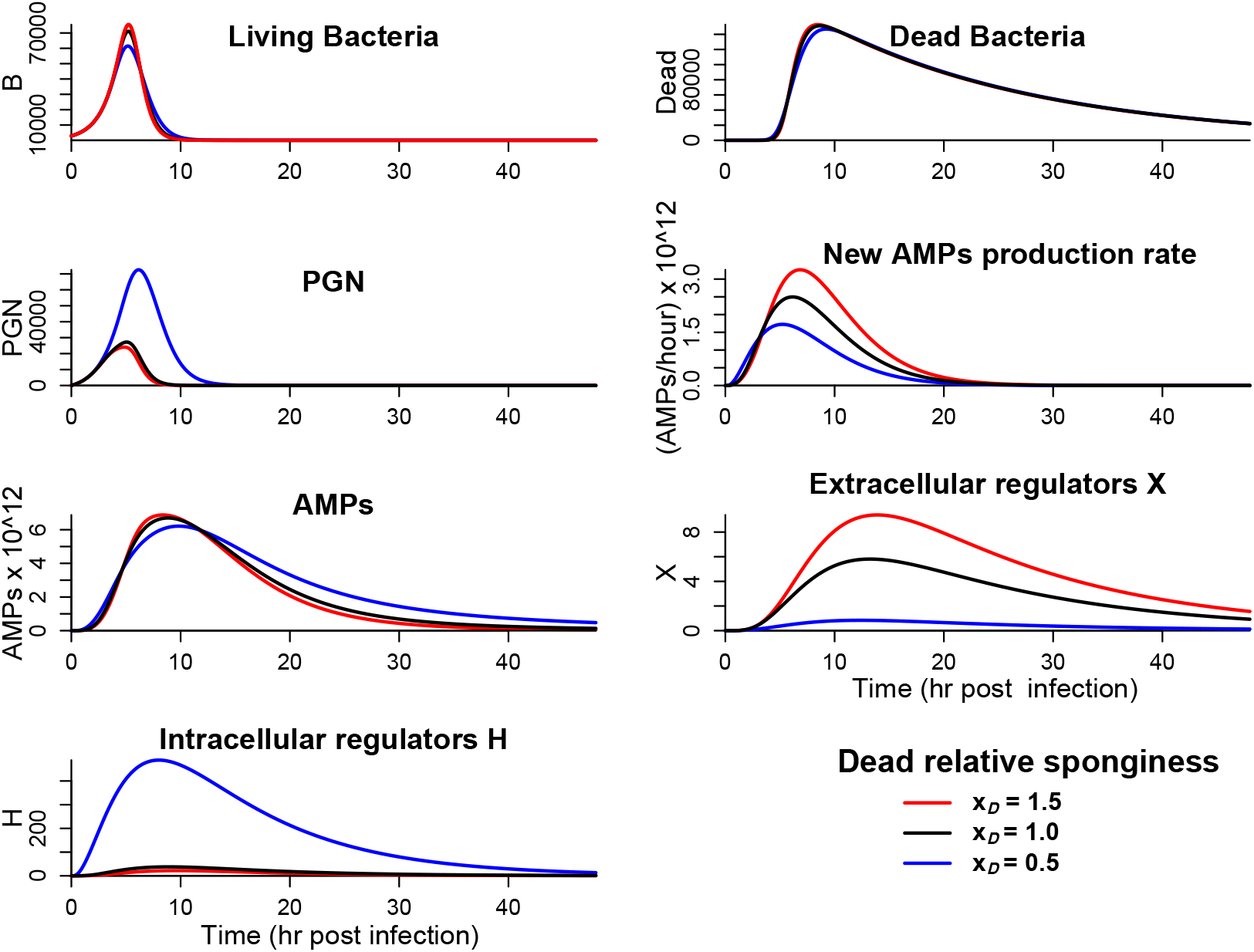
The temporal dynamics of Pareto optimal host immune responses depends on the relative “sponginess” of dead bacteria. Three scenarios with different values of dead-sponginess result in a very similar bacterial damage (within 0.2% from each other), but this is attained through different means. The concentration of intracellular negative regulators is much higher when the sponginess of dead bacterial cells relative to live ones is low, resulting in lower AMP production even though the value of *V*_*A*_ is higher, and lower *X* concentration allowing longer persistence of PGN. The graphs are time plots of (left to right, top to bottom) the numbers of living and dead bacteria, PGN, new AMPs production rate, unattached AMPs, extracellular and intracellular negative regulators. Each panel shows the trajectories for three different Pareto-optimal (*V*_*A*_, *V*_*H*_) values from Fig. 5, with different dead sponginess levels (*x*_*D*_) indicated by colors. Red: (*V*_*A*_ ≈ 1.07 *×* 10^13^, *V*_*H*_ ≈ 13.7); black: (*V*_*A*_ ≈ 1.3 *×* 10^13^, *V*_*H*_ ≈ 38); blue: (*V*_*A*_ ≈ 6.13 *×* 10^13^, *V*_*H*_ ≈ 3.4 *×* 10^3^). Figure made by R script sponge Pareto compare.R

## Discussion

Innate immune effector molecules such as antimicrobial peptides (AMPs) act as “suicide molecules” that are consumed during the immune response. During a controlled infection, many bacteria are killed, generating corpses that can continue to bind and accumulate AMPs, a phenomenon we term “sponginess” of the dead. To explore the physiological consequences of dead bacterial sponginess on infection dynamics, we established a general mechanistic model for pathogens interacting with a stereotypical innate immune response: the systemic response of *Drosophila*. Our model successfully captures essential aspects of host-microbe interactions that occur *in vivo* in *Drosophila*, including the bimodal nature of survival outcomes. Both in experiments (Duneau et al., 2017; Ellner et al., 2021) and in our model, minute variations in infection dose can lead to life-and-death differences in infection outcomes. In agreement with prior literature, our model shows that these outcomes are affected by the strength and ramp-up time of the innate immune response. Additionally, our model suggests that dead bacterial sponginess strongly affects not only the outcomes but also the side effects for the host.

Dead bacterial sponginess has previously been explored *in vitro*, where dead cells can protect their kin from AMPs by sequestering these molecules, resulting in population-level AMP tolerance. However, what has not been explored before now is how this affects within-host infection dynamics and the trade-offs associated with aggressive immune response (Snoussi et al., 2018; Wu and Tan, 2019). During an infection, the degree of dead bacterial sponginess broadens the range of conditions under which the pathogen can overwhelm host defenses. By capturing some of the free AMPs, dead bacterial sponginess shields live pathogens from immune effectors, thus increasing the intensity of immune response required for control. Once the infection is controlled, dead bacterial sponginess switches from being deleterious for the host to being beneficial, through shielding the host from autoimmune damage from free AMPs. The fact that the relative sponginess of the dead, described by the parameter *x*_*D*_ = *c*_*D*_ /*c*_*L*_, is a double-edged sword for the host is confirmed by the sensitivity analysis of our model in Fig. 4. We therefore propose that even moderate levels of sponginess of dead bacteria can be a critical factor driving within-host pathogen dynamics and evolutionary optimization of immune responses. Indeed, the *in vitro* experimental evidence (Savini et al., 2020; Snoussi et al., 2018; Wu and Tan, 2019) suggests that dead sponginess may be far higher than values that already produce major effects in our model.

On the evolutionary time-scale, if the host is regularly exposed to the same level of initial bacterial infection, it is reasonable to expect that the immune response would improve through natural selection to become highly effective for this specific type and level of infection. Absent other selective pressures, one can expect that immune response parameters such as *V*_*A*_ and *V*_*H*_, the maximum prodction rates of AMPs and intracellular negative regulators, would be tuned over generations to minimize a combination of bacterial and AMP-toxicity damage corresponding to specific values of bacterial parameters such as *r*, Λ, and relative dead sponginess *x*_*D*_ . If we neglect the metabolic cost of producing AMPs, the Pareto front analysis in Fig. 5 shows that, even in the case of immune response optimized for each specific dead-sponginess level, higher *x*_*D*_ still results in lower AMP-toxicity damage and/or lower bacterial damage. This holds true despite the fact that the actual strategies of *x*_*D*_ -customized optimal immune response might be quite different, as illustrated for example in the shift to producing mostly intracellular regulators for a lower *x*_*D*_ = 0.5 in Fig. 6. Expecting selection pressures to produce an “optimal” outcome is, of course, naive even with very carefully modeled fitness functions. However, this does not negate the usefulness of considering such (Pareto-)optimal immune response strategies. Rather than predicting a specific evolved immune response, they show the constraints on what evolution could possibly accomplish, a lower bound for the damages that are unavoidably incurred by whatever response actually manifests.

While we have parameterized our model around empirically estimated values wherever possible, empirical data are limited for some aspects. In most cases, this is a function of practical limits in experimentation. Current methods commonly used for quantifying bacterial populations (Dionne et al., 2006; Dostálová et al., 2017; Khalil et al., 2015; Troha and Buchon, 2019) do not separately and simultaneously quantify living and dead bacterial cells over the course of a natural bacterial infection. Furthermore, assays of the AMP sponging by dead bacterial cells have thus far been conducted only *in vitro* (Savini et al., 2020; Snoussi et al., 2018; Wu and Tan, 2019), and their sponginess *in vivo* has never been estimated. The half-life of AMPs in circulation is unknown and are likely to vary among distinct AMPs (Cociancich et al., 1994; Uttenweiler-Joseph et al., 1998). And while some studies have indicated autotoxic damage from AMPs particularly in neurological tissue (Badinloo et al., 2018; Kounatidis et al., 2017; Swanson et al., 2020), the exact amount and tissue-ubiquity of autoimmune damage from AMPs is largely unknown. Finally, the actual production cost of AMPs remains unquantified. Because of these large parameter uncertainties, our evolutionary analysis did not aim to produce quantitative predictions for experimental testing, but is rather a qualitative exploration of the idea that AMP sponging by dead bacteria can have large but entirely different effects on infection and evolutionary time scales.

The present work builds on Ellner et al. (2021) and more recent related studies (Asgari et al., 2023; Hidalgo et al., 2022; Lafont et al., 2022). It introduces several important innovations, including an explicit separation of two types of negative regulators and the first use of a massive multi-hit model in host-microbe interactions. The latter is needed to match biological reality but presents significant analytic and computational challenges, which we handle with the “binning” technique described in Supplement *§*S4.2. We provide some analytic results on the stationary distribution of AMP hits in a bacterial population subject to a constant density of AMPs and under the assumption that the death occurs after (Λ + 1) hits only; see Supplement *§*S6. Given the estimated mortality function, which rapidly transitions from low to high risk of death before dividing, it may be adequate to assume that bacteria are killed by some critical number of hits and unaffected by any smaller number.

Our model reaffirms the crucial importance of rapidly eliminating an infection that has succeeded in becoming systemic (Fig. 4). The most influential parameters for all measures of disease impact are the time-constant for induction of immune response, the rate at which AMPs are produced once the response is induced and the rate at which AMPs binds to living pathogens, and the number of AMP “hits” needed to kill instantly (*τ*_*U*_, *V*_*A*_, *c*_*L*_ and Λ respectively). Parameters affecting the shape of bacterial mortality as a function of hit number are less important (*µ*_0_, *θ*), which is fortunate because those are difficult parameters to experimentally quantify.

In our study of immune system evolution, we have considered only two of many potentially important defense costs (e.g., we considered auto-toxicity of AMPs but not the cost of producing those AMPs), and only two potentially evolvable parameters in the host’s immune response (*V*_*A*_ and *V*_*H*_). Taking account of both production costs and autoimmune damage is an important area for future study, although a proper exploration will require accurate empirical estimates of AMP production costs that do not currently exist. This will also allow for a more careful consideration of the role played by other evolved host parameters; e.g., the mechanisms for clearing out bacterial corpses – see Supplement *§*S8.

Another direction for future work will be to explore the evolutionary optimality of the overall immune system architecture. For example, how do hosts benefit from producing both intracellular negative regulators that block immune signaling as well as extracellular negative regulators that degrade the immune stimulus? And why do negative regulators and activating surveillence molecules exist as multigene families of closely related paralogs (Dziarski, 2004)?

Here we have considered a single infection, but a natural, free-living host is likely to face multiple infections, and its altered immunological state after fighting off one infection may change the starting conditions for a second that arises before the host re-equilibrates. Evolution of the immune response therefore may depend on how frequently such rapid-succession infections occur. Moreover, pathogens vary in their growth rates, tolerance of AMPs, AMP sponginess, and detectability by the host immune system. The consequences of a response strategy will depend on which pathogens a host is exposed to during its lifetime, increasing the challenge of devising an immune system that can cope with a large suite of encountered pathogens.

Our results demonstrate that dead bacterial sponginess can strongly affect within-host infection dynamics and evolutionary optimization of the immune system. Some empirical estimates of *x*_*D*_ in recent papers suggest that dead bacterial cells may be considerably more spongy than we have assumed here (Savini et al., 2020; Snoussi et al., 2018; Wu and Tan, 2019), suggesting that the real impacts of shielding by the dead may actually be much larger than our theoretical predictions. This shielding may also contribute to additional aspects of infection, including the empirically observed chronic infection sustained by surviving hosts (Chambers et al., 2019). Altogether, our results demonstrate that dead bacterial sponginess is an overlooked but potentially crucial element of host-microbe interactions. Elucidating its full role will require an iterative combination of mathematical modeling and empirical experimentation.

More broadly, our results highlight a general (and underappreciated) principle: “killed” enemies are often not inert, but can persist on infection time-scales as structured biochemical particles that bind, sequester, or otherwise neutralize host effectors, thereby reshaping the effective strength, and side-effects, of host defense. For AMPs specifically, multiple studies show that dead bacteria can absorb and retain large numbers of peptide molecules, protecting surviving cells and increasing population-level tolerance to AMPs. When incorporated into within-host dynamics, this “corpse sink/decoy” effect changes not only whether infections clear, but also the shape of the fundamental trade-off between pathogen control and immunopathology, thereby altering which immune activation/shutdown strategies can be favored by natural selection as environments, pathogens, or host physiology vary. This extends beyond *Drosophila*: by changing within-host pathogen loads and infection duration, corpse-mediated shielding can ultimately affect transmission and virulence evolution, and thus selection on hosts and pathogens. Finally, this perspective suggests two general evolutionary hypotheses: pathogens might be selected to enhance corpse persistence or effector binding as a kin-benefitting “public good,” while hosts might be selected to invest in more rapid clearance/processing of microbial debris (analogous in spirit to the strong selection for corpse clearance in other immune contexts) to prevent dead pathogens from interfering with host defenses. The potentially large effects of AMP sponging by the dead raise the question of whether killed pathogens might also benefit their living kin in other ways, in the context of both innate and adaptive immune responses, further generating selection pressures on the host for investing in clearing mechanisms even if they are costly.

## Acknowledgements

This project was supported by NIH (R01 AI141385 to BPL, R01 AI148529 and AI148541 to NB, and R01 AI143704 to TD), NSF (IOS 2024252 to NB, DMS 2111522 to AV, DEB 1933497 to SPE and G. Hooker), AFOSR (FA9550-22-1-0528 to AV), USDA-NIFA (AFRI grant 2021-67015-35235 to S. McArt, C.Myers, and SPE, as part of the USDA-NSF-NIH-UKRI-BSF-NSFC Ecology and Evolution of Infectious Diseases program) and the Royal Society (Wolfson Visiting Fellowship to AV). MIK is supported by a Fleming Postdoctoral Fellowship. Any opinions, findings, conclusions, or recommendations expressed in this publication are those of the author(s) and do not necessarily represent official views of the funding organizations. We are grateful to four reviewers for comments that substantially improved the presentation.

## Statement of Authorship

NB, TD, SPE, BPL and AV contributed to conceptualizing the study, model development, discussion and interpretation of results, and writing the manuscript. TD designed and supervised experiments that were performed by MK. SPE and AV did the coding and math. SPE wrote most of the first manuscript draft and all authors besides MK contributed to revisions.

## Data and Code Availability

All original data and all computer scripts necessary to replicate the results are available from https://doi.org/10.5281/zenodo.18481456 Ellner et al. (2026).

## Online Supplement

### S1 Detailed model development

#### S1.1 Sensing and responding to an infection

Presence of bacteria is signaled by an increase in PGN *G*, which is perceived by the host as an indication of infection and stimulates an immune response. We assume that the rate of PGN shedding by living cells is proportional to the cell division rate (Nigro et al., 2008). Because cell death caused by AMPs is associated with lysis, we assume that the PGN from a killed cell becomes immediately accessible for recognition by the host at the moment of cell death. In our focal experimental system, PGN triggers the Imd signaling pathway (and to a much lesser extent the Toll pathway (Hixson et al., 2024; Troha et al., 2018), which we ignore for simplicity). The result of the signaling cascade is nuclear localization of the NF-*κ*B transcription factor Relish (modeled as production of active Relish) and binding to promoter sites of genes that express host immune proteins. This includes genes encoding antimicrobial peptides, as well as negative regulators, peptidoglycan sensors, and Relish itself (Buchon N, 2014). PGN is probably always present at some very low level, coming from the host’s microbiota, but we ignore this because it is far smaller than the amount produced quickly at the onset of a potentially harmful infection.

An essential feature of the immune response is the time delay between signal and response, which can determine the outcome of the infection (Duneau et al., 2017; Ellner et al., 2021). We model the delay as follows. A constant fraction of bound nuclear promoter sites (resulting from some constant PGN concentration) would produce after several hours a corresponding steady rate of AMP production. We let 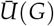 denote that production rate as a fraction of the maximum rate when all promoter sites are bound. A detailed model of the Imd signaling cascade (see *§ S2 Input-output description of the initial cascade*) suggests that the relationship between PGN and steady-state bound nuclear Relish is approximately a Monod curve. We assume that a sufficiently high PGN concentration would result in nearly all promoter sites being bound, while extracellular negative regulators *H* decrease the sensitivity to *G*, so we specify 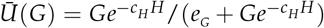 for some constants *e*_*G*_ and *c*_*H*_ .

Up-regulation of AMP production is the result of multiple steps and we model it as such:

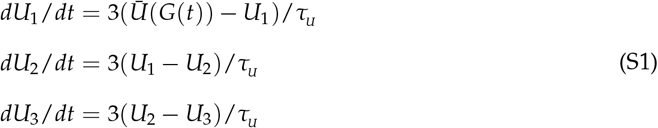

Given enough time, if *G*(*t*) is constant then 𝒰_1_, *𝒰*_2_, and *𝒰U*_3_ all converge to 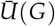. Our use of three steps was motivated by considering that ramping up of AMP production is mainly delayed by transcription, translation, and secretion, while the signaling cascade and promoter binding are much faster. We assume that the host dies, and production of AMPs and other gene products ceases, when the total bacterial population *B* reaches a threshold value Ω (called the BLUD, bacterial load upon death, by Duneau et al. (2017)). AMP production rate is given by *V*_*A*_*𝒰U*_3_*W*(*B*/Ω) where *V*_*A*_ is the maximum possible rate, and *W* is a function that equals 1 when *B «* Ω and quickly but smoothly shuts down production when *B*/Ω increases towards 1 (see SI Fig. S1). The chain of state variables *U*_*j*_ produces a gamma-distributed delay between a brief spike in *G* and consequent up-regulation *U*_3_, with *τ*_*U*_ as the mean of the distribution. A single step model produced unrealistically fast control of infections relative to bacterial population increase (i.e., the ratio between peak and initial bacterial populations was much smaller than observed experimentally (Duneau et al., 2017)), while the three-step model produces more realistic dynamics of controlled infections (see *§ Results*, Fig. 1 in the main text) and has a mechanistic interpretation.

**Figure S1.**
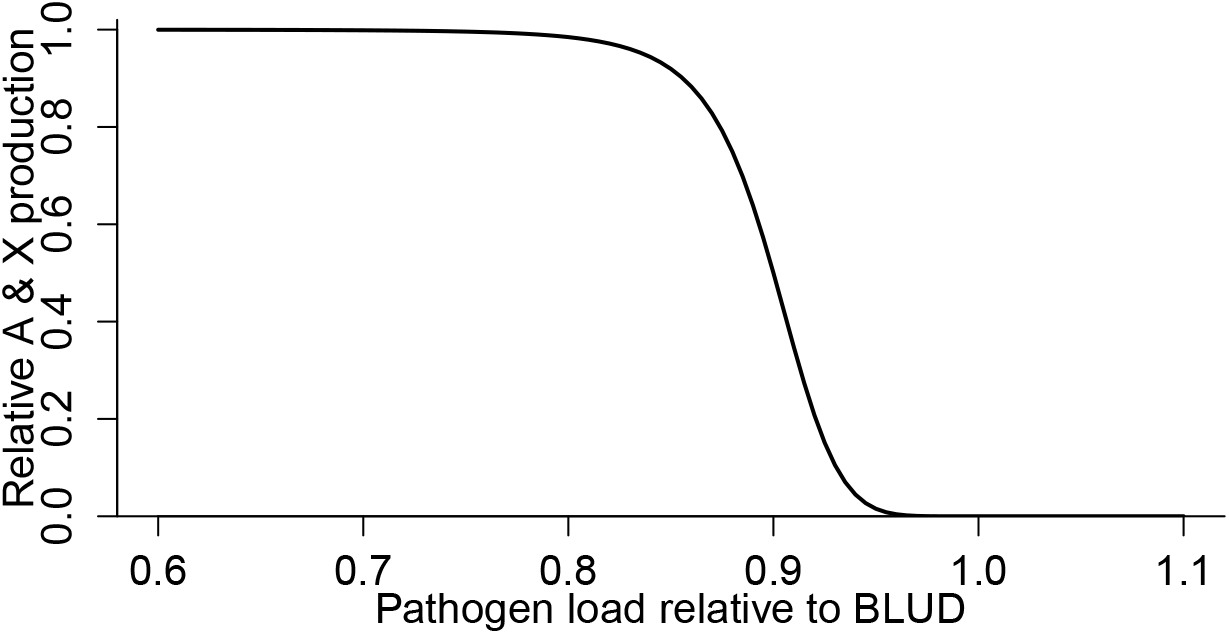
The host “well being” function *W* showing how AMP and extracellular regulator *X* production decrease when the pathogen load *B* approaches the threshold level Ω. We used the function *W*(*x*) = (1 − *x*)^6^)/(0.1^6^ + (1 − *x*)^6^) for 0 ≤ *x* ≤ 1 and *W*(*x*) = 0 for *x >* 1 (meaning that the host is dead because bacterial load is above Ω). Figure made by `Multihit_utilities.R.`

AMP molecules are lost through binding to living and dead bacteria, by natural degradation with first-order loss constant *δ*_*A*_, and through degradation by proteases that are produced by the bacteria. As in Ellner et al. (2021) we make the quasi-steady-state assumption that protease concentration is proportional to bacterial abundance. This implies an AMP loss rate *mAB* representing the effect of proteases. We assume that binding of AMPs to host tissue, which causes AMP auto-toxicity, never becomes saturated and is therefore modeled as a first-order process and combined into the loss constant *δ*_*A*_ .

Binding of nuclear Relish also up-regulates production of negative regulators. Intracellular regulators *H* decrease the sensitivity to PGN, as described above. Extracellular regulators degrade PGN and thus dampen further stimulation of the immune response after an infection has been controlled. Although there are multiple such regulators (e.g., families of PGRPs with amidase activity (Bischoff et al., 2006; Zaidman-Rémy et al., 2006)), we combine them into a single variable *X* with production rate *V*_*X*_*𝒰U*_3_. We assume first-order kinetics for *X* degrading PGN, creating a PGN loss term *ρ*_*G*_ *XG*. We assume that *X* and *H* are only lost through natural degradation with decay constants *δ*_*X*_ and *δ*_*H*_ respectively.

#### S1.2 Bacterial population dynamics

We model AMP-induced bacterial mortality as a multi-step process, building on past models for killing by antibiotics (Abel Zur Wiesch et al., 2015; Clarelli et al., 2020; Hemez et al., 2022). This results in a delayed and gradual response of the bacterial mortality rate to dynamic changes in AMP concentration, whereas classic pharmacokinetic-pharmacodynamic models (Minichmayr et al., 2022; Witzany et al., 2023) posit an immediate response dependent on AMP concentration and cell state (e.g., active versus resting). Gradual response due to cumulative binding of additional AMPs is biologically more realistic when AMP is produced at physiologically reasonable rates (in contrast to a therapeutic antibiotic that is instantaneously applied at an overwhelming concentration). This lagged response is important because it magnifies the number of cells that must be killed to control an infection, and may also be important for evolutionary optimization of the host immune response if bacteria cause damage to the host before the infection is brought under control (see *§ Results* in the main text).

We assume that bacterial mortality is zero in the absence of AMPs, while each additional “hit” (bound AMP molecule) produces an increase in mortality rate, described by a smooth function *µ*(*x*) giving the instantaneous mortality rate for a cell with *x* hits. We let Λ denote the maximum number of hits that a live cell can sustain – when a cell with Λ hits acquires one more, it immediately dies. The live bacterial population thus consists of sub-populations *B*_0_, …, *B*_Λ_ with the subscript encoding the number of bound AMP molecules. The total population of live bacteria is *B* = *B*_0_ + … + *B*_Λ_. We assume that all sub-populations have the same maximum division rate *r* and overall carrying capacity *K*. We also assume that AMP molecules bind to all live cells at the same rate *c*_*L*_ *A*, because the number of possible binding sites is very high relative to Λ. This property of AMP binding differs from the situation with many antibiotics, where binding sites can be saturated at clinically relevant concentrations (Abel Zur Wiesch et al., 2015).

Consider first the gains and losses of cells in *B*_*j*_ due to cell divisions. We assume that when a cell divides, each bound AMP molecule is equally likely to go to either daughter, independent of the fate of other AMP molecules. In the calculations below, we consider that division by such a cell eliminates it from *B*_*j*_. On the other hand, a daughter cell in *B*_*j*_ can be produced by division of cells in compartments *B*_*j*_, *B*_*j*+1_, … *B*_Λ_.

When a cell in *B*_*j*+*m*_ divides (with *m* ≥ 0), we can arbitrarily label one of them as the “first daughter”. If the first daughter has *j* AMPs, this creates a daughter in *B*_*j*_. If the first daughter has *m* AMPs, this also creates a daughter in *B*_*j*_ because the second daughter necessarily has *j* AMPs.

Thus, for each cell division in *B*_*k*_ with *k* = *j* + *m*, the expected number of daughters in *B*_*j*_ is

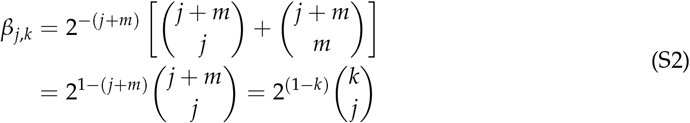

Combining losses and gains to *B*_*j*_ through cell divisions with those due to AMP binding, the living bacterial population dynamics are

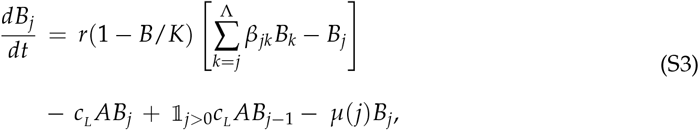

*j* = 0, …, Λ. The first term on the right hand side is changes due to cell divisions (with the factor 1 − *B*/*K* reflecting resource limitations on cell division rate), the second is “promotions” out of *B*_*j*_ into *B*_*j*+1_ due to AMP binding, and the next is promotions into *B*_*j*_ from *B*_*j*−1_ due to AMP binding (for *j >* 0), and the final term is mortality.

As we lack direct information on hit-dependent mortality *µ*(*j*), we specified a flexible functional form and estimated its parameters by optimizing model fit to experimental data and data summaries (see *§*S5). We initially assumed a product of three terms: a gradual power-law increase with per-capita mortality proportional to *j*^*θ*^, *θ >* 0, a step-like increase at some critical number of hits, and a rapid increase near Λ hits such that cells with Λ + 1 hits die instantly, proportional to 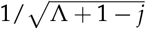. However, the step increase was invariably estimated to be zero so we removed it, giving 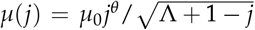 . The rapid increase near *j* = Λ also had little effect because the fitted *µ*_0_ and *θ* values produce rapid death at *j* well below Λ (see Fig. S8), but we retain it to ensure that all living bacteria have *j* ≤ Λ hits.

Analysis of (S3) shows that the AMP concentration needed to prevent bacterial population growth (if *A* is held constant) is at most *r*(1 + Λ)/*c*_*L*_ ; this bound represents the situation where bacterial mortality is zero until a cell acquires Λ + 1 hits and immediately dies. Because the estimated *µ*(*j*) increases steeply at *j* = Λ/2, *r*(1 + Λ/2)/*c*_*L*_ is a reasonable estimate of the AMP concentration at which killing by AMPs exactly balances cell divisions.

Because Λ is very large (*O*(10^7^) hits, see *§*S3), eqn. (S3) yields an extremely high-dimensional system of stiff coupled nonlinear ODEs. For that reason, we used a coarsened (“bin-based”) approach to obtain accurate numerical model solutions; see *§*S4.2 for details.

#### S1.3 AMPs sponging by the dead

We assume the simplest possible model for AMP binding to a killed cell, using first-order reaction kinetics. We assume that bacterial cells are very large targets for AMP binding, and that if one dead bacterial cell breaks into several fragments, the cumulative binding potential of the fragments remains high enough that there is no effect on overall AMP binding potential in the system. The degradation rate of “corpses” (and their corresponding fragments) and their elimination from the system is given by *δ*_*D*_ . The model for the number of dead bacteria *D* is then

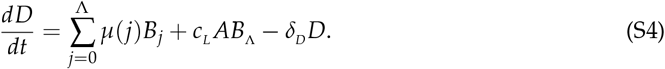

#### S1.4 Rescaling the model

The process models developed in the previous sections combine to produce the overall model. Before studying the model, we can reduce the number of free parameters by rescaling variables. We scale time so that one unit is the bacterial doubling time at the maximum possible population growth rate *r*, resulting in *r* = ln(2) ≈ 0.69. For typical bacterial opportunistic pathogens, the doubling time *in vivo* is on the order of an hour (Gibson et al., 2018; Haugan et al., 2018), so we can informally think of time as measured in hours. We scale *G* so that one bacterial division produces one unit of *G* (*γ* = 1), and then scale *X* so that the rate constant for degradation of PGN is *ρ*_*G*_ = 1. Then *γ*_*D*_ represents the PGN production from killing one bacterium, relative to the amount produced when one living cell divides. We scale *H* so that *c*_*H*_ = 1. However, we keep live and dead bacteria dimensional (bacteria per host), for comparison with experimental results. We cannot rescale AMPs usefully because the exact integer number of hits and maximum number Λ affect the distribution of hits in daughter cells. These scalings produce the complete model shown in the main text. Ranges of values for model parameters based on our focal experimental system are discussed in *§*S3 and *§*S5.

### S2 Input-output description of the initial cascade

In this section we derive an input-output description 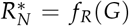 of the Imd signaling cascade, where 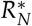 is the amount of bound nuclear Relish. The description is based on assuming quasi-equilibrium of the initial cascade relative to the current value of PGN, *G*. We are mostly interested in the possible qualitative shapes of this function so the dynamic model in the main text can use a simple but realistic parametric equation *f*_*R*_ instead of including additional dynamic equations for the signaling cascade that would introduce many additional model parameters.

Our model of the cascade is diagrammed in Fig. S2. This is essentially the same in as in Ellner et al. (2021), except that we omit the intracellular positive regulators that maintain the supply of Relish needed for the cascade. Here we assume for simplicity that those regulators do their job perfectly, and as a result the parameters of the cascade (i.e., all of the rate constants in the reaction kinetics) are constant. Our dynamic model in the main text includes intracellular negative regulators, denoted *H*. Their effect is modeled as a reduction in the sensitivity to PGN, so that 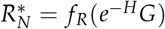 when units of *H* are scaled appropriately.

**Figure S2.**
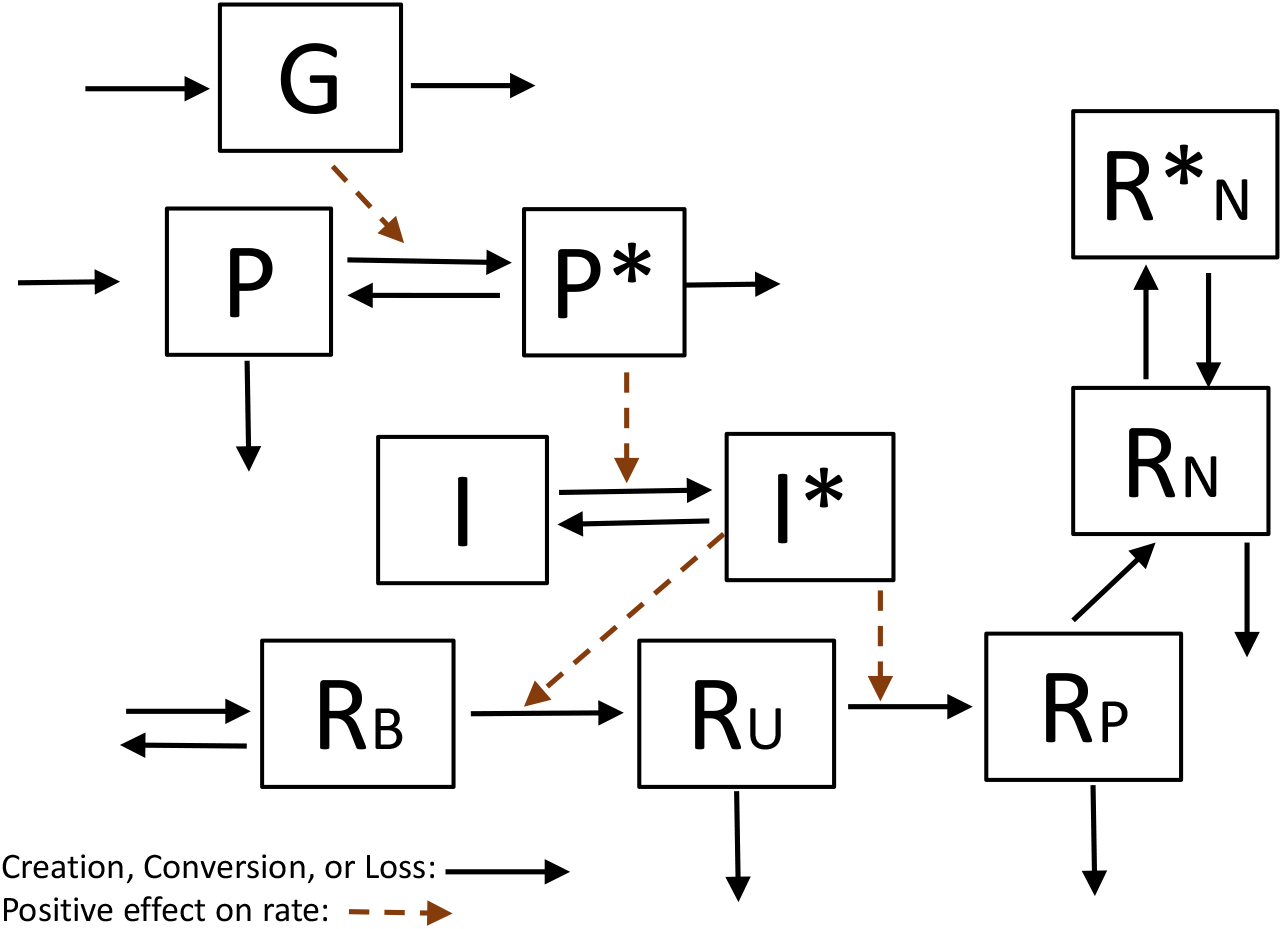
Our simplified model for the initial signaling cascade from PGN to bound nuclear Relish, in the fork of the IMd pathway going through PGRP-LC. Solid black arrows indicate flows. Dashed red lines indicate positive effects of a variable’s concentration on the rate of a reaction arrows The state variables are *G*: peptidoglycans exterior to the cell (moles/l); *P, P*^*^: unbound and bound PGRP-LC (moles); *I, I*^*^: free and recruited IMD (moles); *R*_*𝒰*_, *R*_*B*_, *R*_*P*_: forms of Relish outside the nucleus; 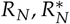: unbound and bound Relish in the nucleus. *R*_*B*_ is the native cytoplasmic version of the protein, having an intact self-negative-regulatory domain. *R*_*𝒰*_ is cleaved – the negative domain is removed – but it is still unphosphorylated and therefore inactive. *R*_*P*_ is both cleaved and phosphorylated and therefore is active.

For convenience we omit below brackets [•] denoting concentration.

*The first step* is dimerization of PGRP-LC (*P*) through binding to PGN (*G*). On the fast time scale of these chemical processes we can assume that the total amount of PGRP-LC is a constant *P*_*T*_. The reaction diagram

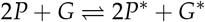

gives the kinetic equation

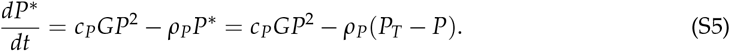

In the loss term, *ρ*_*P*_ is the detachment rate of dimers, the concentration of dimers is *P*^*^ /2, and each dimer detachment event produces two unattached *P* molecules.

We assume that *G*^*^ is negligibly small compared to *G*. We define *e*_*P*_(*H*) = *c*_*P*_/*ρ*_*P*_, the ratio between the forward and backward rate constants (*e* for “efficiency”). Then solving the quadratic (S5) for *P* we get the steady states

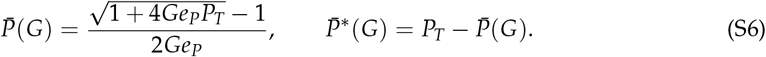

This expression already rules out finding the overall cascade steady-state analytically. The Taylor series about *G* = 0 (using Maxima script PGN.max) is 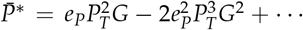, increasing and concave down. That remains true for all *G*: the first derivative in *G* is positive (converging to 0 as *G* → ∞), the second derivative is negative. Fig. S3 shows 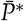 as a function of *G* for *P*_*T*_ = *e*_*P*_ = 1. The limit as *G* → ∞ is 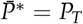, i.e. all PGRP-LC is in the dimerized state.

**Figure S3.**
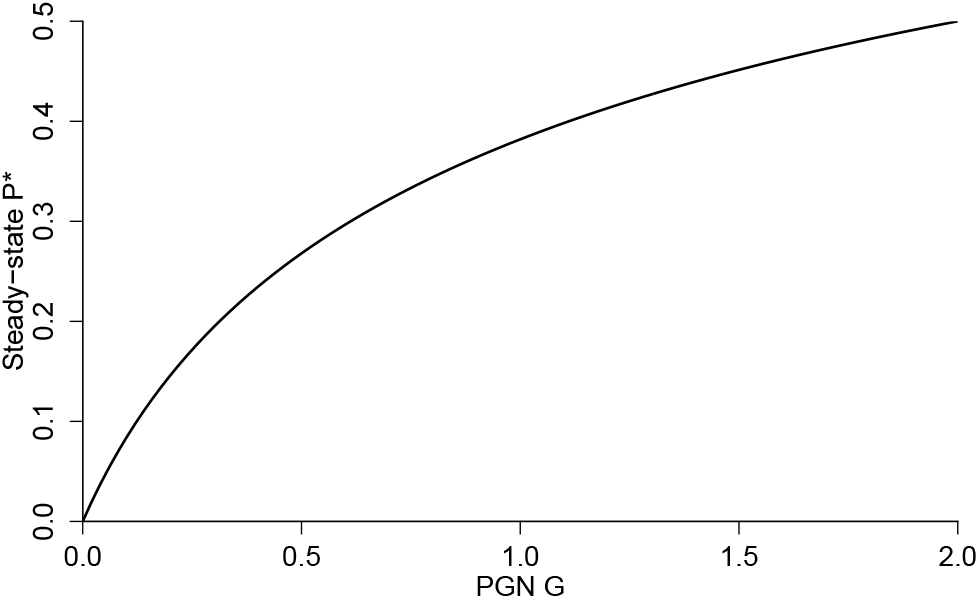
Graph of steady-state 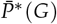 as a function of *G*, for *P*_*T*_ = *e*_*P*_ = 1.

*The second step* is recruitment (activation) of Imd, *I* ;::: *I*^*^ with *I* + *I*^*^ ≡ *I*_*T*_, giving kinetic equation

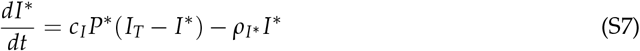

The steady-state value is Michaelis-Menten,

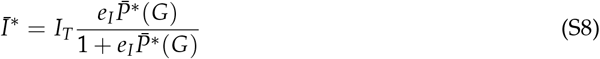

where *e*_*I*_ = *c*_*I*_/*ρ*_*I*_. As a function of *G* (with 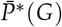) as found above) this has the same qualitative shape as 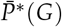, increasing and concave down (why: if functions *f* and *g* are both increasing and concave down, it’s simple calculus to prove that their composition is increasing and concave down).

*The third step* is the one we omit in our kinetic model: recruited Imd catalyzing activation of kinase and caspase. Instead, we assume that activated kinase and caspase are both proportional to *I*^*^. The reaction kinetics for kinase and caspase both have the same form as Imd, so the actual quasi-equilibrium relationships are both Michaelis-Menten functions of *I*^*^. Thus, we tacitly assume that concentrations of activated caspase and kinase are small enough that saturation as a function of *I*^*^ can be ignored.

*The fourth step* is Relish outside the nucleus:

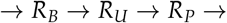

with *R*_*B*_ → *R*_*𝒰U*_ and *R*_*𝒰U*_ → *R*_*P*_ catalyzed by *I*^*^. It is reasonable to assume that *R*_*𝒰U*_ goes “immediately” to *R*_*P*_, i.e. the *R*_*𝒰U*_ → *R*_*P*_ step is very fast. The leftmost → is production of *R*_*B*_; the rightmost → is one-way transport of *R*_*P*_ into the nucleus (where it becomes *R*_*N*_). Previously we collapsed this step by omitting *R*_*𝒰U*_, but that may be problematic for our present purposes if it changes the functional form of the steady-state rate of transport into the nucleus.

Binding of nuclear Relish leads to up-regulation of *R*_*B*_ production. The kinetic equations are

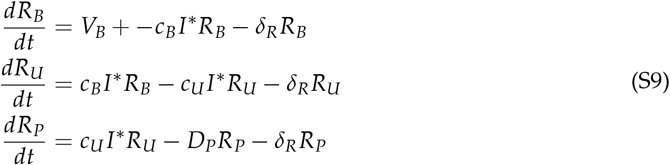

The total amount of cytoplasmic Relish results from the balance between production of *R*_*B*_, natural degradation, and transport of *R*_*P*_ into the nucleus. Setting all derivatives in (S9) to zero and solving, we get that the steady-state of *R*_*P*_ is

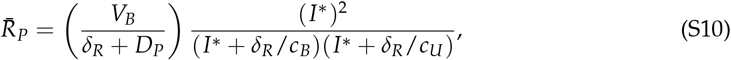

in other words

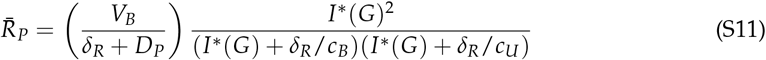

where *I*^*^ (*G*) is given by eqns. (S8) and (S6). *f*_*R*_ is thus an increasing, saturating function of *G* that is quadratic (concave up) near *G* = 0.

The next steps are movement of *R*_*P*_ into the nucleus (becoming *R*_*N*_), and binding/unbinding of *R*_*N*_ with promoter sites:

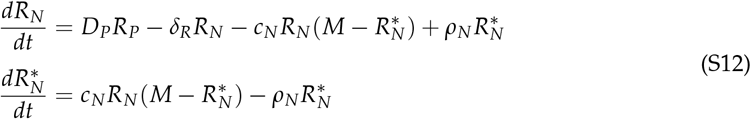

where *M* is the total number of promoter sites at which *R*_*N*_ can bind. At steady state the second line equals 0, hence the last two terms in the first line add to zero, implying 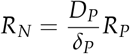 state. Solving the second line gives 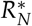 in terms of *R*_*N*_. Putting those together, at steady state

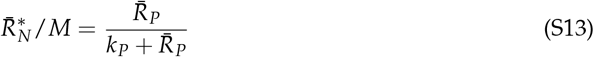

where *k*_*p*_ = (*ρ*_*N*_*δ*_*P*_)/(*c*_*N*_ *D*_*P*_).

We now want to explore the possible qualitative shapes of 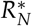 as a function of *G*. As a first step, we can reduce the parameter count by expressing the result in terms of parameter combinations, as follows.

First, *e*_*P*_ only appears in the product *Ge*_*P*_, so for examining the qualitative shape we can take *Ge*_*P*_ as the independent variable. Equivalently, without loss of generality we can assume *e*_*P*_ = 1.

When (S8) is substituted into (S11), we can divide numerator and denominator by *I*_*T*_ and the only effect is to change the value of the *δ*/*c* constants in the denominator of the second factor. We can therefore set *I*_*T*_ = 1 in (S8) and make the constants in the denominator of the second factor our basic parameters.

Second, when (S11) is substituted into (S13), we can divide numerator and denominator by the first right-hand-side factor in (S11), and this only changes the value of *k*_*p*_. After these steps *k*_*p*_ is a combination of parameters, including many that do not appear anywhere else in the formula for 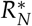 . We can therefore take *k*_*p*_ as a basic model parameter.

With these simplifications, the input-output relationship is given by combining (S13) with (S6) for 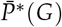 with *e*_*P*_ = 1, and (S8) with *I*_*T*_ = 1, giving

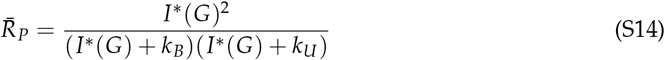

The free parameters are thus *P*_*T*_, *e*_*I*_, *k*_*B*_, *k*_*𝒰U*_ and *k*_*P*_.

Generating curves of 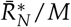 versus *G* with randomly drawn parameters, we find that that except for very small values of *G* (i.e., values such that most promoter sites are unbound), the curve is qualitatively like a Michaelis-Menten curve (Fig. S4). In the main text, we therefore assume that the fraction of bound promoter sites is given by a Michaelis-Menten function of *G*.

**Figure S4.**
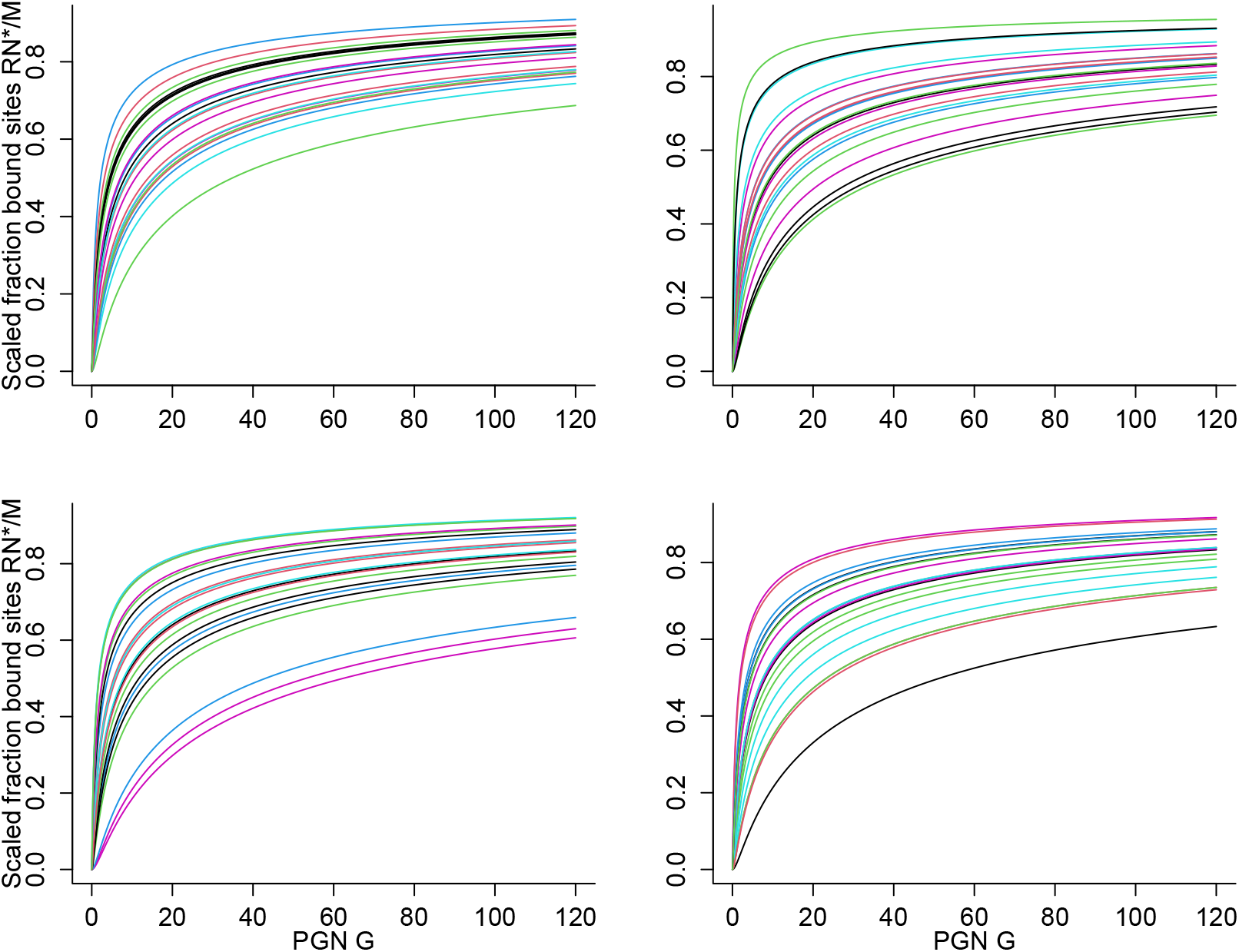
Graphs of 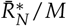 versus *G* with randomly drawn parameters. Each panel shows the result of random parameter draws. All curves are scaled relative to their maximum value as *G* → ∞. Parameter sets were generated as *iid* lognormal perturbations of base values (*P*_*T*_ = 1, *e*_*I*_ = 2, *k*_*B*_ = 1, *k*_*U*_ = 1, *k*_*P*_ = 1) e.g., the random value of *e*_*I*_ in each parameter set was generated as 2*e*^*Z*^ where *Z* has a Normal(0,1) distribution. Figure made by `R_script_CascadeFunction.R`

### S3 Parameter ranges

Here we explain the rationale for our choice of baseline parameter values for the parameters whose values were not derived by model calibration (*§*S5).

1. Ω, the BLUD (bacterial load upon death). A typical value is 5 *×* 10^6^ bacteria per fly (Duneau et al., 2017).
2. *K*, the “carrying capacity” represents total consumption of the host and so is much larger than Ω. So long as *K »* Ω the particular value has no effect on any of the properties we are studying, so we arbitrarily set *K* = 1000 *×* Ω.
3. *τ*_*𝒰*_ . Lemaitre et al. (1997) found that transcription of mRNAs encoding AMPs was generally detectable but low at 1 hour after infection with a variety of bacteria, and strong and increasing at 3hr and maximal by 6 hr. In our model, transcription rate increases from zero to maximal, in response to a constant AMP concentration, in proportion to 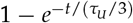. Values of *τ*_*𝒰*_ /3 between 2 and 3 hours produce up-regulation in our model that align with the qualitative results of (Lemaitre et al., 1997). We therefore chose *τ*_*𝒰*_ = 7.5 hours as the baseline value for our model. After calibration of other model parameters (as described below) this value produced a good match to two observed features of the dynamics in our focal experimental system (Duneau et al., 2017). First, there is significant AMP up-regulation detectable by 4 hours after infection with several thousand bacteria. Second, there was significant continued increase after 4 hours, such that AMP levels sufficient to bring an infection under control (when that did occur) were reached only after ≈ 7 hours following inoculation with the pathogen, and the bacteria population continued to grow approximately exponentially until close to that time.
4. *γ*_*D*_ is the total PGN released as the result of a cell death (we assume that this release is instantaneous), relative to that released when one living cell divides. The dead do have immunostimulatory effects (Hixson et al., 2024; Troha et al., 2018). But in experiments where equal numbers of living and dead bacteria were injected into different hosts (mosquitoes, *Aedes aegypti*), living bacteria induced a far stronger immune response (Hixson et al., 2024). Translating this empirical finding into a value for *γ*_*D*_ is challenging because, in an experiment, the immune response to living cells creates dead cells in the same host, and because the living potentially increase in numbers over the course of infection. A further complication is the nonlinear and time-delayed relationship between PGN concentration and immune response. We therefore considered that *γ*_*D*_ was likely to be somewhere between 0.01 and 1, and applying our geometric mean principle (described in *§ Choice of default parameter values* of the main text) we adopted *γ*_*D*_ = 0.1 as the baseline value. This means that most PGN stimulation of the immune system in our model comes from living bacteria, unless the dead greatly outnumber the living. We further note that, because *γ*_*D*_ has very low elasticity for all of the response measures that we considered (too small for it to be included in Fig. 4), mis-estimation of this parameter would have very little effect on model predictions.
5. *m*. A bacterial cell removes AMP molecules from circulation two different ways: by absorbing AMPs (acquiring “hits”) and by producing protease, an enzyme that degrades AMPs but is not consumed in the process. The model assumes a constant ratio between the number of bacteria and the protease concentration. The ratio between AMP elimination rate by proteases and AMP elimination rate through absorption, at any given moment, is given by the parameter *m*. Knocking out protease production has only a small effect on bacterial growth in *Drosophila* infected with *Serratia marcescens* bacteria (Flyg and Xanthopoulos (1983) and our own unpublished experiments). We therefore set *m* = 0.1. This is perhaps small enough to simply ignore, but we retain *m* in the model because larger values might apply in other systems.
6. *x*_*D*_ gives the “sponginess” of dead bacteria relative to live bacteria, so we are interested in exploring the range *c*_*D*_ = *O*(1). We chose default value *x*_*D*_ = 1 because cell death does not alter the chemical properties relevant to AMP absorption, but we consider that larger values are worth exploring. For example, the dead may lack active defenses that the living have.
7. *δ*_*A*_ . As in Ellner et al. (2021), we consider that AMPs are stable on the time scale of an infection (i.e., the ≈ 0.5 − 1 day required for clearing an infection or host death). Documented half-lives for different AMPs vary from hours to a week (Cociancich et al., 1994; Uttenweiler-Joseph et al., 1998). If we take the range of half-lives to be approximately 1/8 day to 8 days, the geometric mean principle gives 1 day as a reasonable half-life. As baseline value we use *δ*_*A*_ = 0.03, giving AMP a half-life of 23.1 hours.
8. *δ*_*X*_ . The extracellular negative regulators are enzymes, similar to lysozymes, that degrade PGN. Lysozyme half-lives are highly variable, ranging from 1 to 50 hours (Inoue and Rechsteiner, 1994*a*). On the geometric mean principle we assume a half-life of ≈ 7 hours, giving the default value *δ*_*X*_ = 0.1. The value of *δ*_*X*_ turns out to be fairly unimportant for the properties we consider, because once *X* has done its job of degrading PGN, it matters very little how much longer it persists in the fly.
9. *δ*_*H*_ . Many different types of protein are actually involved in intracellular negative regulation, each of which will have different kinetics, and little is known about their half-lives. A typical half-life for a protein would be several hours (i.e., neither minutes nor days) (Eden et al., 2011; Inoue and Rechsteiner, 1994*b*). We set *δ*_*H*_ equal to *δ*_*X*_ so that the two components of the immune shut-down process operate on the same time scale.
10. *δ*_*G*_ . PGN is also fairly stable on the time scale of infection success or failure (Jutras et al., 2019). Exact values of these parameters are not important for the dynamics of model simulations, so long as they are not far from *δ*_*A*_ and *δ*_*X*_ . We chose *δ*_*G*_ = 0.03 giving a half-life of ≈ 24 hours.
11. *δ*_*D*_ . The loss of dead bacteria and thus of their ability to sponge AMPs is affected both by chemical decay and by active clearance of dead cells by the host through phagocytosis and/or filtering. Our system is modeled on adult *Drosophila*, in which phagocytes are rare and easily saturated (Guillou et al., 2016), so we chose a relatively low value of *δ*_*D*_ = 0.05 giving dead cells (or more precisely for our model, the AMP-sponging capacity of dead cells) a half-life of 14 hours. The situation would be very different in *Drosophila* larvae, or in other insect species with more macrophages and high filtration capacity, but we believe that it is the correct choice for the model system represented by our default parameters. Nevertheless, to explore the robustness of our conclusions for species with more active clearance of dead cells, we replicate our main analyses for a scenario where dead cells are cleared 10*×* faster, *δ*_*D*_ = 0.5, but are more AMP-spongy than living cells (which is in line with the experimental evidence (Savini et al., 2020; Snoussi et al., 2018)); see *§*S8 below.
12. *e*_*G*_ . This parameter controls how strongly the host responds to the presence of a small bacterial population. With *B « K* bacteria present, *dG*/*dt* = *rB* − *δ*_*G*_ *G* so the value after 1 hour starting from *G*(0) = 0 is *G*(1) ≈ *rB* ≈ 0.7*B* because degradation within an hour is negligibly small. When we allowed *e*_*G*_ to be a fitted parameter in model calibrations (described below), the best fits all had implausibly small values, *e*_*G*_ *«* 1 or smaller, implying a fully activated immune response to the PGN produced by one bacterium in an hour. As a minimal lower bound, we assume that the PGN produced by 10 bacteria in an hour should elicit a weak response so as not to over-react to a harmless bacterium “passing through”, suggesting *e*_*G*_ ≥ 100. Assuming larger values up to *e*_*G*_ = 300 had little effect on calibrated values of other parameters, and no visible effects on model dynamics with best-fit parameters, so we kept *e*_*G*_ = 100 as our default value.
13. The experimental results on shielding by heat-killed bacteria (Main Text, Methods and Materials, *Experimental demonstration of shielding by heat-killed bacteria*) provide a rough estimate for Λ, the maximum possible number of hits on a living bacterium. For these calculations, we assume that living and heat-killed cells are more or less equally “spongy”, because heat-killed cells (unlike AMP-killed cells) still have intact membranes, at least initially. In the experiment, the presence of 10^7^ heat-killed bacterial cells had a very small effect on mortality of living cells but 10^8^ heat-killed cells greatly reduced the living cell mortality. Thus, we infer that 10^8^ heat-killed cells were enough to absorb, within well under an hour, an appreciable fraction of the AMP molecules initially present and thereby shield the living cells (absorption in the same trial by the 7 *×* 10^6^ initially living cells is small enough to ignore). So if 10^8^ killed cells together acquired *O*(10^14^) to *O*(10^15^) hits (and thus, a non-negligible fraction of the 10^15^ AMP molecules present) each heat-killed cell would have acquired *O*(10^6^) to *O*(10^7^) hits, and each initially living cell would have acquired about the same. The instantly lethal dose Λ therefore must be substantially larger than this number of hits. This suggests that the highly lethal hit burden Λ is *O*(10^7^) or possibly larger, which is consistent with previous estimates in the literature (Roversi et al., 2014; Savini et al., 2020). As an upper bound, if all of the 10^15^ initially present AMP molecules were absorbed by the 7 *×* 10^6^ initially living cells, each cell would have about 1.4 *×* 10^8^ hits on average. Consequently, that number of hits must be high enough to entail substantial mortality. Thus, Λ is unlikely to be much larger than 10^8^. We chose Λ = 2 *×* 10^7^ as our default value.

By similar arguments, we attempted to roughly estimate the AMP binding coefficient *c*_*L*_ from the same set of experimental results. However, we found that this required a number of questionable assumptions and still left nearly hundred-fold uncertainty about the value. We therefore put *c*_*L*_ among the parameters to be estimated by model calibration, as described in *§*S5.

### S4 Numerical solution of the model

#### S4.1 Making sure that extinct is forever

Regardless of the AMP concentration, the number of bacteria in the model cannot fall to exactly zero; at best, bacteria decline exponentially toward zero. However, this type of spurious persistence can allow a modeled bacterial population to regrow (in model solutions) even after the host’s immune response has reduced them to far less than one living cell, with bacteria eventually becoming so numerous as to kill the host. This triumphant return can occur because the negative regulators *X* and *H* might still be at appreciable concentrations in the host, preventing an effective immune response, while previously produced AMPs would have been absorbed by living or dead bacteria.

In order to prevent this unrealistic behavior, we imposed a “quasi-extinction” boundary at low bacteria densities. In ecological terminology, we added an artificial strong Allee effect at low cell counts. Specifically, the cell division rate *r*(1 − **B**/*K*) was multiplied by **B**^4^/(3^4^ + **B**^4^) where **B** is the total number of bacteria. All other per-capita rates of the bacteria, including mortality, were multiplied by **B**^4^/(1 + **B**^4^); see Fig. S5. The nonlinear reduction in cell division rate, relative to other rates, implies that at sufficiently low cell counts, divisions are dominated by mortality. This behavior is biologically realistic, as very small bacteria populations are likely to be held in check by processes not included in our model, such as phagocytosis.

**Figure S5.**
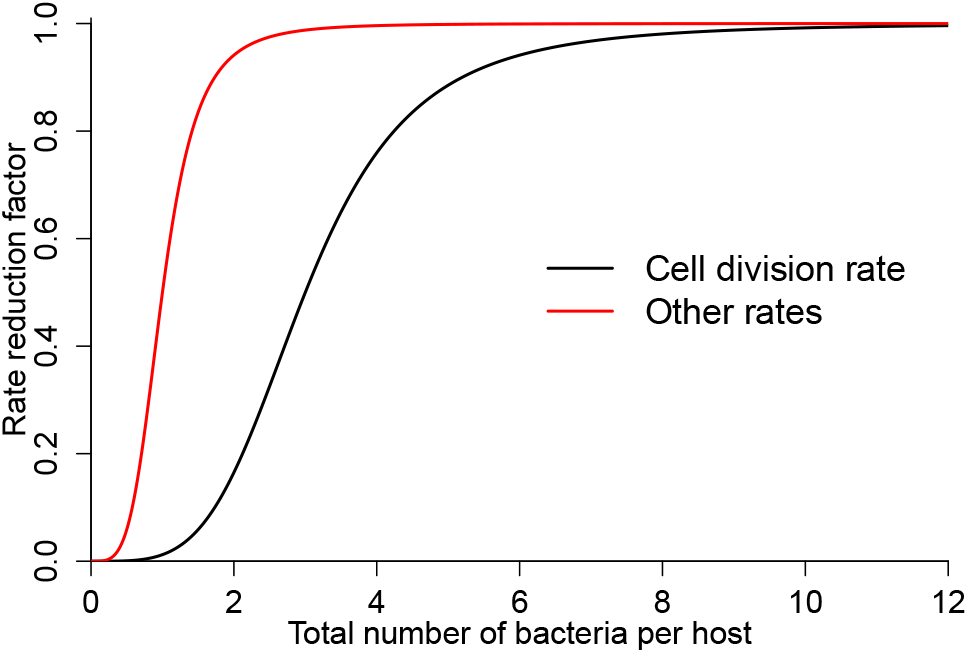
The rate-reduction functions used to impose an extinction boundary at very low bacteria cell counts. When the total number of bacteria cells in the host falls below 10, cell division rate begins to decline. Other rates also decline, but only when the cell count is below 1.Figure made by `Bin_scheme.R.`

The smooth decrease in other rates is a device to prevent negative bacteria abundances in numerical solutions of the model. The more common device of using log-transformed state variables fails for our model because very low values of state variables (especially, bacteria being driven to minute cell counts) produce numerical overflow in the calculating the logarithm of a very small number, crashing solver routines.

In our bacterial population equations with these modifications, bacteria in the absence of a host immune response would be a one-dimensional ODE with a stable equilibrium at **B** = 0, and an unstable equilibrium at **B** = *O*(1) cells in a host. With initial conditions above the unstable equilibrium the bacteria will increase to their carrying capacity. Initial conditions below the unstable equilibrium lead to extinction of the bacteria, with the extinction rate slowing as **B** = 0 is approached due to the declining per-capita mortality rate. In the full model, a bacteria population driven below the unstable equilibrium by the host immune response will inevitably decrease to zero, while the dynamics for **B** = 10 or more (which is 2 order of magnitude below their experimental initial abundance) are unaffected by the modifications.

#### S4.2 Approximation by binning

In the model bacteria, are classified by AMP “hits”, the number of bound AMP molecules running from 0 to Λ where Λ is very large (our default is Λ = 2 *×* 10^7^). The model therefore cannot be integrated directly. Instead, we use a low-dimensional approximation in which bacteria are combined into “bins” representing an interval of AMP hit numbers.

Recall that when a cell divides, hits are allocated independently, with equal probability, to the two daughters. This implies that the fraction of divisions, by parents with *k* hits, that produce a daughter with *j* ≤ *k* hits, is

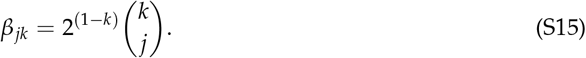

Let *z* denote “hit fraction” *k*/Λ where *k* is the absolute number of hits. Consider a cell with hit fraction *z* that divides. Label one daughter as the “first”. The number of hits in the first daughter is Binomial with parameters *N* = *z*Λ and *p* = 0.5. The mean number of hits is *Np* = *z*Λ/2, corresponding to hit fraction *z*/2 in the daughter. But the variance in the number of hits is is *Np*(1 − *p*) = *z*Λ/4, and so the variance of the resulting daughter *z* values is *z*/(4Λ).

Higher Λ gives a tighter distribution, hence a larger number of bins is needed to approximate the distribution. Moreover, the variance is lower for lower *z*. This creates problems for solving the model using even-size bins. Suppose cells are classified into 1 + *s* bins based on hit number,

where *λ* = Λ/*s* is the “bin width”. Bin 0 consists of cells with exactly zero hits; bin 1 ≤ *£* ≤ *s* consists of cells with (*£* − 1)*λ* + 1 to *£λ* hits. When a cell in bin *£* divides, the expected number of daughters sent to bin *m* would be the average, across the hit numbers in bin *£*, of the expected number of daughters that lie in bin *m* according to (S15).

To illustrate the effects of small variance at low hit numbers, consider for example Λ = 10^5^ with 200 even-size bins. Bin 0 consists of individuals with 0 hits, Bin *j >* 0 consists of individuals with 500(*j* − 1) + 1 to 500*j* hits. The computed allocation matrix bins the 8 lowest bins is

|  |  |  |  |  |  |  |  |  |
| --- | --- | --- | --- | --- | --- | --- | --- | --- |
| FROM: | 0 | 1-500 | 501-1000 | 1001-1500 | 1501-2000 | 2001-2500 | 2501-3000 | 3001-3500 |
| TO: |  |  |  |  |  |  |  |  |
| 0 | 2 | 0 | 0.00 | 0.00 | 0.00 | 0.00 | 0.00 | 0.00 |
| 1-500 | 0 | 2 | 1.95 | 0.05 | 0.00 | 0.00 | 0.00 | 0.00 |
| 501-1000 | 0 | 0 | 0.05 | 1.95 | 1.93 | 0.07 | 0.00 | 0.00 |
| 1001-1500 | 0 | 0 | 0.00 | 0.00 | 0.07 | 1.93 | 1.91 | 0.09 |
| 1501-2000 | 0 | 0 | 0.00 | 0.00 | 0.00 | 0.00 | 0.09 | 1.91 |
| 2001-2500 | 0 | 0 | 0.00 | 0.00 | 0.00 | 0.00 | 0.00 | 0.00 |
| 2501-3000 | 0 | 0 | 0.00 | 0.00 | 0.00 | 0.00 | 0.00 | 0.00 |
| 3001-3500 | 0 | 0 | 0.00 | 0.00 | 0.00 | 0.00 | 0.00 | 0.00 |

Here, reading down column *j* gives the bin distribution of daughters from bin-*j* parents. Bin 0 is exact. But bin-1 and bin-2 parents both produce nearly 100% bin-1 daughters, and bin-3 and bin-4 parents both produce daughters that are almost entirely in bin 2. It continues like that. With evenly spaced bins, pairs of adjacent bins are assigned nearly identical daughter distributions, both with nearly zero variance.

This problem can be reduced by using narrower bins at small *z*. In order to approximately represent the variance of the allocation distribution, each bin’s width should be comparable to (and ideally smaller than) the standard deviation of the allocation distribution for daughter cells being added to that bin.

Let 0 = *x*(0) *< x*(1) *< x*(2) *<* … *x*(*N*) = 1 denote the bin boundaries on *z* scale, and 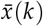 the midpoint of the *k*^*th*^ bin. We can heuristically think of *x* as real-valued. The typical hit number in bin *k* is 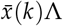, so incoming daughter cells were produced by parents with roughly 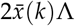 hits. The hit distribution in their daughters has variance 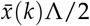 in hits, and therefore standard deviation 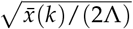 on the *z* scale. We therefore want

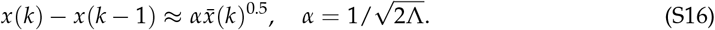

Again, let us treat *k* as a continuous variable *u*, and approximate the last equation as

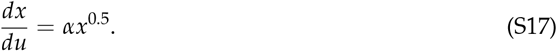

The solution is 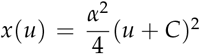 where *C* is a constant of integration. We need (very roughly speaking) *x*(*u*) → 0 as *u* → 0, so *C* ≈ 0. Returning to discrete bin boundaries *x*(*k*), we conclude that *x*(*k*) should increase quadratically with *k*, and hence bin widths increase roughly linearly with *k* (because (*k* + 1)^2^ − *k*^2^ = 2*k* + 1).

Because the variance becomes very low at low hit numbers, we used a hybrid scheme in which the first *m* + 1 bins consist of cells with exactly 0, 1, 2, …, *m* hits, and thereafter we have heterogeneous bins whose widths increase linearly. Function `make_limits` in script `Multihit_funs_binned` implements the hybrid scheme. With Λ = 10^6^ (a 10-fold increase from the equal-bins example), *m* = 4 and 120 heterogeneous-size bins, the allocation matrix for bins the 8 lowest bins is

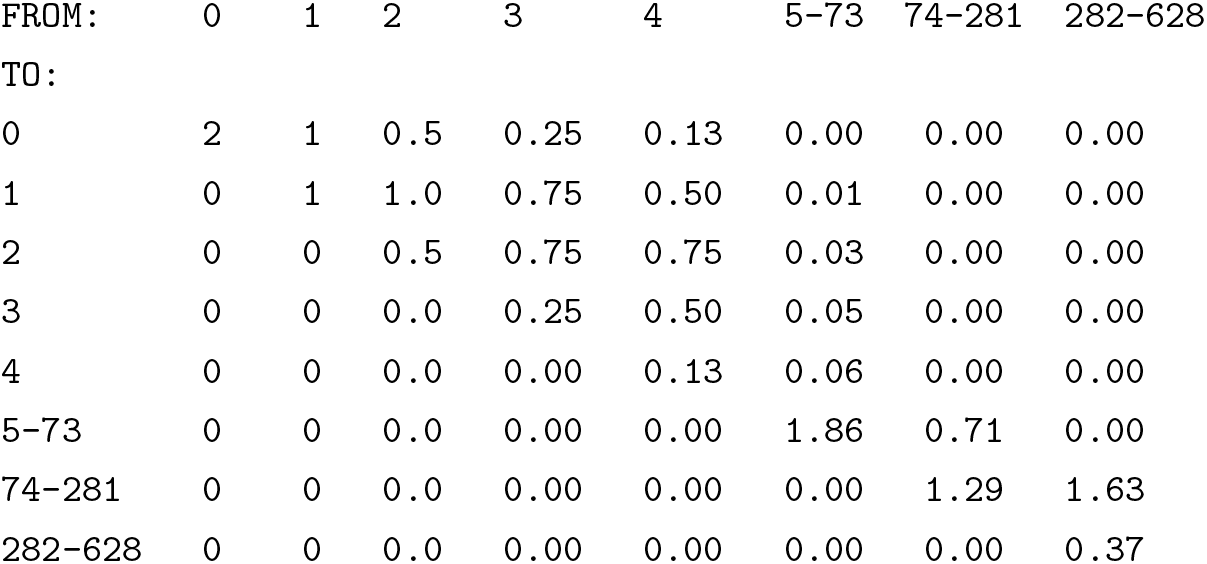

| FROM: | 0 | 1 | 2 | 3 | 4 | 5-73 | 74-281 | 282-628 |
| --- | --- | --- | --- | --- | --- | --- | --- | --- |
| TO: |  |  |  |  |  |  |  |  |
| 0 | 2 | 1 | 0.5 | 0.25 | 0.13 | 0.00 | 0.00 | 0.00 |
| 1 | 0 | 1 | 1.0 | 0.75 | 0.50 | 0.01 | 0.00 | 0.00 |
| 2 | 0 | 0 | 0.5 | 0.75 | 0.75 | 0.03 | 0.00 | 0.00 |
| 3 | 0 | 0 | 0.0 | 0.25 | 0.50 | 0.05 | 0.00 | 0.00 |
| 4 | 0 | 0 | 0.0 | 0.00 | 0.13 | 0.06 | 0.00 | 0.00 |
| 5-73 | 0 | 0 | 0.0 | 0.00 | 0.00 | 1.86 | 0.71 | 0.00 |
| 74-281 | 0 | 0 | 0.0 | 0.00 | 0.00 | 0.00 | 1.29 | 1.63 |
| 282-628 | 0 | 0 | 0.0 | 0.00 | 0.00 | 0.00 | 0.00 | 0.37 |

The first 5 columns are exact, and the others approximate the variability among daughter cells better than with even-size bins.

For most of our calculations, we used *m* = 10 and *n* = 120 so that the bacteria population is approximated by 131 bins (see *§*S4.4 below).

#### S4.3 ODEs with unequal bin widths

The equations for living and dead bacteria populations need to account for the heterogeneous hit-number bins with nonconstant width. Let **M** be the daugher-allocation matrix with variable bin widths, *w*_*j*_ the width of bin *j* (the number of distinct hit numbers in the bin), *µ*_*k*_ the instantaneous mortality rate of a cell with exactly *k* hits, and 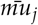 the average of *µ*_*k*_ over the hit numbers in bin *j*.

Indexing on bins and entries of **M** requires some care because the first bin corresponds to zero hits, not one. We therefore let bin indices, and the subscripts on matrix entries **M**_*jk*_, run from 0 to *S*, where bin *S* has Λ as its right endpoint.

Let *cA* be the rate at which living cells aquire hits when AMP concentration is *A*. If AMP concentration is constant at *A*, then it takes a cell *w*_*j*_/(*cA*) time units, on average, to go from entering bin *j* to leaving bin *j* in the absence of mortality. The advection rate out of cell *j* should therefore be set to *cA*/*w*_*j*_.

Combining losses and gains to *B*_*j*_ (the number of living cells in bin *j*) through cell divisions, AMP binding, and mortality, the living bacterial population dynamics are implemented numerically as

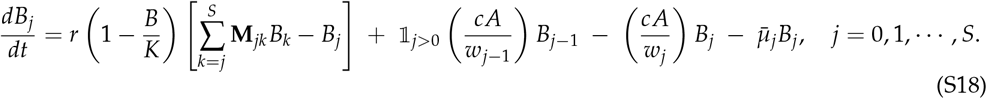

The first term on the right hand side is changes due to cell divisions, the next two are “promotions” into and out of *B*_*j*_ due to AMP binding, and the last is mortality. There are no promotions into *B*_0_, and cells “promoted” out of bin *S* are dead because Λ + 1 is an instantly lethal dose. Note that the promotion terms can be grouped for *j >* 0 as

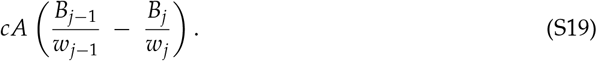

The term in parentheses is an “upwind” discrete concentration gradient, so our numerical scheme makes “physical” sense.

The dynamics of dead cells are then approximated numerically as

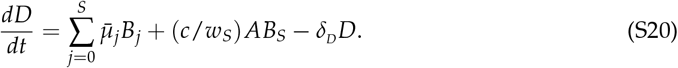

The first term is mortality losses from within bins, the second is instantaneous death upon acquiring Λ + 1 hits, and the third is loss through natural degradation.

#### S4.4 How many bins is enough?

Choosing the number of bins is a trade-off between accuracy (more bins) and computing speed (fewer bins). Our computing bottlenecks in this study were parameter optimizations, first for estimating parameters through trajectory matching, and then for evolutionary optimization (see main text *§ Materials and Methods*). By trial and error, we found that for parameter values producing either controlled and uncontrolled infections at initial bacterial concentrations close to those where bimodal outcomes were observed by Duneau et al. (2017), using more than *m* = 10 (11 exact bins) had no detectable effect on the dynamics of controlled infections, large increases above *n* = 120 heterogeneous bins had minute effects on solutions but appreciable effects on solution time, and decreasing *n* much below 120 had visible effects on solution accuracy.

Fig. S6ABC shows the dynamics of total bacteria population, AMP concentration, and immune-upregulation *U*_3_ with *n* = 120 and *n* = 240 heterogeneous bins, for a controlled infection using our default parameter values. Small differences are apparent at the peaks of each, but the differences are very small (1.26%, 1%, and 0.23% of the average for bacteria, AMPs, and *U*_3_, respectively). In contrast, increasing from *n* = 120 to *n* = 240 heterogeneous bins increases the time required to calculate the plotted solutions by a factor of 2.8. We expect the greatest effects of numerical errors for initial conditions at the split between controlled and lethal infections. Fig. S6D shows the maximum controlled bacteria population in numerical model solutions for different numbers of heterogeneous bins. Bin number does affect the precise value, but again the effect is very small: for *n* between 80 and 280, the value changes by less than 1%.

**Figure S6.**
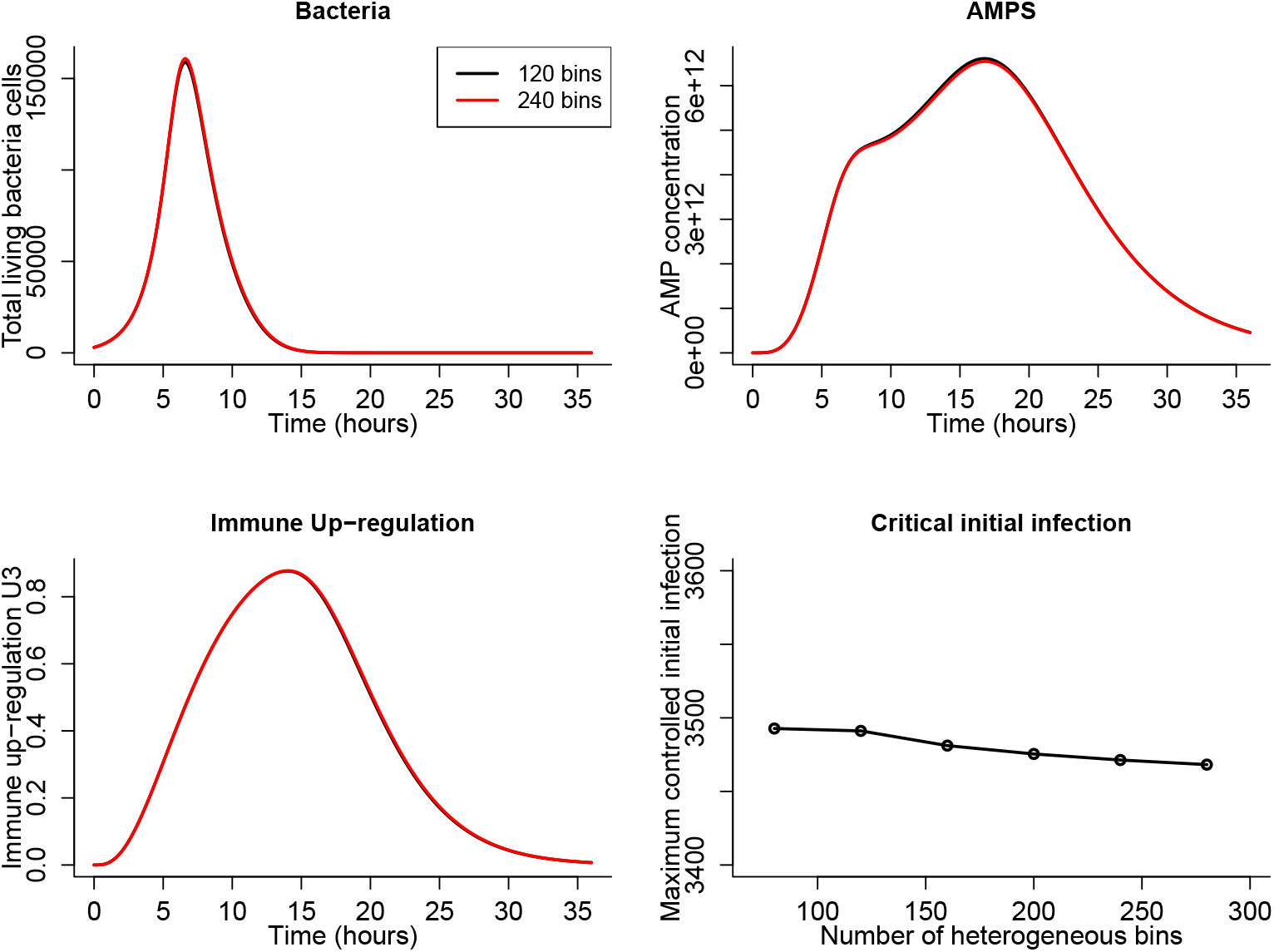
A,B,C: Numerical solutions of the model with default parameter values, *m* = 10 (11 exact bins) and *n* = 120 or *n* = 140 heterogeneous bins. D: Maximum initial infection (number of bacteria present at *t* = 0, all with zero AMP hits) such that the bacteria population is driven to extinction by the host immune response, rather than growing to the carrying capacity *K* and killing the host. Figure made by `Compare_matrix_size.R.`

### S5 Parameter estimates from model calibration

Here we explain how the parameters *V*_*A*_, *V*_*X*_, *V*_*H*_, *c*_*L*_ and the parameters in the mortality function 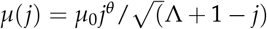 were estimating by minimizing an error function quantifying the overall discrepancies between model predictions and the three performance targets described in the main text. The descriptions here provide additional details on how the error terms were calculated from model simulations.

The first target was the observed survival rate in the experiments with heat-killed bacteria (Main Text, Methods and Materials, *Experimental demonstration of shielding by heat-killed bacteria*), in the treatments with 10^7^ and 10^8^ bacteria. As noted in the Fig. 1, the observed survivals in the other treatments are interpreted as a very small number of bacteria (one in 10^5^ − 10^6^) that are resistant to AMPs, which are omitted from our model because they would have no effect on the dynamics in the situations that we consider in this paper. Model predictions were generated by initializing the living and dead bacterial populations to the intended densities at the start of the experiment for each treatment, with all living bacteria having zero AMP hits, and eliminating any host immune response because these experiments were done *in vitro*. The error function term *e*_1_ from this target was the sum of absolute differences between the natural logs of observed and predicted survival rates in the two treatments.

The second target was a pair of stylized trajectories of total living bacteria population dynamics (Fig. S7) representing typical behavior seen in the experiments in Duneau et al. (2017).

**Figure S7.**
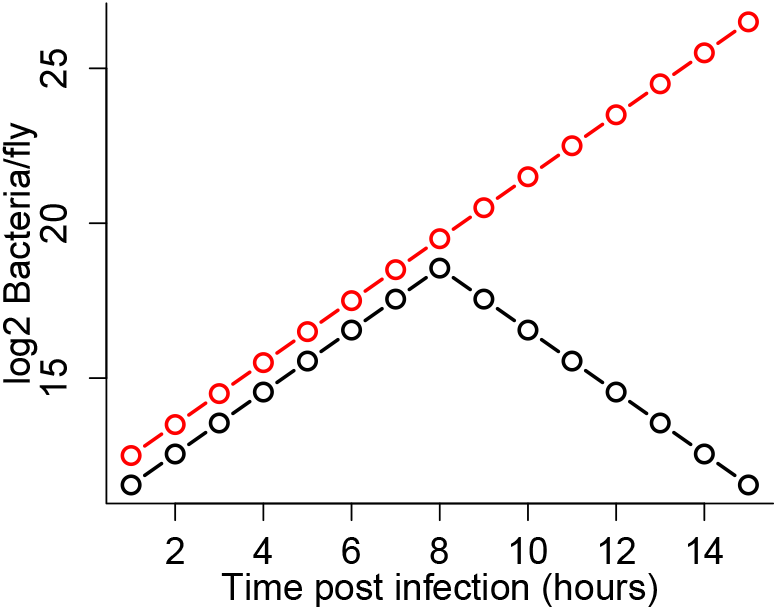
Stylized bacterial population trajectories, typical of those seen in Duneau et al. (2017), exhibiting divergent outcomes as a function of the total number of bacteria in the host when the presence of a pathogen is detected and the IMD signaling pathway is triggered. Figure made by script `Figure_stylized_split.R`

Model predictions for each trajectory were generated by initializing the living bacterial population to the first “data point” exactly with all living bacteria having zero AMP hits, and all other state variables set to zero (in particular, no PGN, AMPs, or dead bacteria, and no up-regulation of AMP production). The error function term *e*_2_ from this target was the sum over the two trajectories of the mean absolute difference between the natural logs of observed and predicted total living bacteria.

The third target was the property that AMP concentrations decrease monotonically after reaching their maximum during the course of a controlled infection. The error function term *e*_3_ from this target was the fraction of time that *dA*/*dt* was positive after the time when AMP concentration reached its maximum in a simulation of the model from *t* = 0 to 72 hours. The simulation was initiated as for the second target, with 2^11.5^ living bacteria present at *t* = 0, which is very slightly below the initial bacterial population for the lower (controlled infection) trajectory in Fig. S7.

The weights attached to the error function terms were adjusted by trial-and-error so that parameter values that were optimal or near-optimal resulted in model simulations matching all three targets fairly well, rather than matching two very well and not coming at all close to the third. The overall error function that parameter values were chosen to minimize was **J** = 4*e*_1_ + *e*_2_ + 25*e*_3_. The large coefficient on the *e*_3_ term makes it act as penalty term ensuring that optimal parameters give *e*_3_ = 0, meaning that AMP concentrations increase to their maximum and then decline monotonically to zero in the course of a controlled infection.

For numerical evaluation of the objective function **J**, the mortality function was re-parameterized in terms of the mortality rate *µ*_1/2_ at *j* = Λ/2 hits and the exponent *θ >* 0,

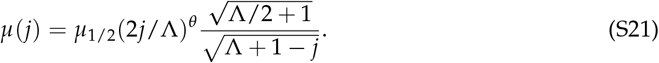

Numerical minimization of **J** was done in stages. Beginning from an initial guess at the optimal parameters, nearby parameter vectors were generated at random by adding Gaussian random values with mean 0 and standard deviation 0.1 to the log of the initial guess and exponentiating. Any time the value of **J** at the new random parameter vector was smaller than the value at the initial guess, the random parameter vector replaced the initial guess as the base point for generating random parameter vectors. This was repeated for 250 draws of random parameter values. The resulting “rough guess” was used as the starting point for minimization by the Nelder-Mead simplex method using the optim function in R, with a maximum of 1000 iterations. The resulting “best” parameters were used as the starting point for another Nelder-Mead minimization; and this was repeated for up to 5 minimizations.

We found that the exponent *θ* was problematic: the final value after multiple Nelder-Mead optimizations was always very close to the initial guess. Moreover, repeating the optimization with *θ* “frozen” at its final value invariably found parameter values giving a smaller value of **J** — which is mathematically impossible, if the numerically recovered minima really were minima. This difficulty arose, we believe, because any changes in *θ* without a precisely compensating change in *µ*_1/2_ caused large increases in **J**, so that any virtually change in *θ* was unsuccessful. To cope with this, we carried out a series of optimizations in which a value of *θ* was assumed; see Fig. S8. The final value of **J** was relatively insensitive to the assumed value of *θ*, with shallow and nearly identical local minima around *θ* = 3 and *θ* = 9. We considered the sudden mortality increases for *θ* ≥ 9 to be unlikely: the odds of death before dividing go from 50:50 at ≈ 1.25 *×* 10^7^ hits, to 10:1 at ≈ 1.6 *×* 10^7^ hits. We therefore focused on *θ* ≈ 3. When penalties were added to constrain 2 ≤ *θ* ≤ 4, optimization resulted in *θ* ≈ 3. We therefore fixed *θ* = 3 and used optimization as described above to select the default values of the other calibrated parameters.

**Figure S8.**
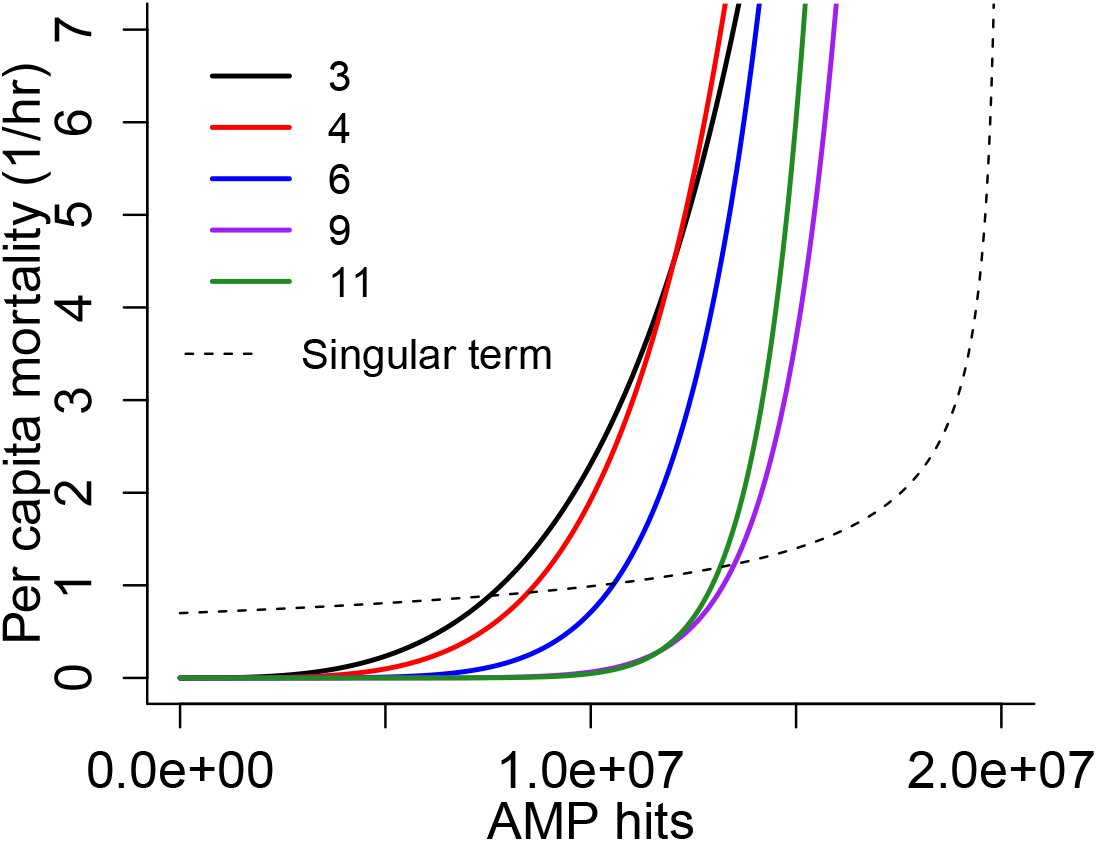
Best-fit mortality curves for different initial values of the exponent *θ*; final best-fit values of *θ* were all within a few percent of the initial values. The dashed curve shows the singular term in 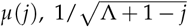 using our default parameter value Λ = 2 *×* 10^7^. Figure made by script Compare `mortality_fits.R`

We remind readers again that the currently available data are not adequate to estimate all model parameters with confidence (see main text *§ Discussion*). Our goal is to demonstrate the potential importance of killed bacteria, in a model with parameter values that are consistent with all available data and estimated as well as we can.

### S6 The AMP concentration required for pathogen control

In this section we derive an upper limit on the AMP concentration required from control of the pathogen, that is, the value of *α* = *cA* (in the unscaled model) such that the long-term bacteria population growth rate is 0. The upper limit, given by *α* = *r*(1 + Λ), is derived by simplifying the model in two ways: we assume *K* = ∞, allowing unlimited exponential growth of the bacteria population if AMPs are absent, and we assume that mortality is zero until AMP concentration reaches the instantly lethal pathogen concentration Λ + 1. While seemingly extreme, this approximation is not entirely irrelevant because the fitted mortality rate function rises quickly in a fairly narrow range, from values such that a cell is very likely to divide before dying (i.e., mortality *«* 0.69/*hr*) to values such that a cell is very likely to die before dividing. Replacing Λ with the location of that range in the results below should give a good approximation for the actual model’s behavior.

We let *x*_*j*_ denote population fractions *x*_*j*_ = *B*_*j*_/**B**, where **B** = ∑_*i*_ *B*_*i*_. Let ***β*** be the matrix of *β*_*jk*_ values, and **G** be the matrix with -1 on the diagonal and 1 on the first subdiagonal. Then the model with *K* = ∞ can be written as

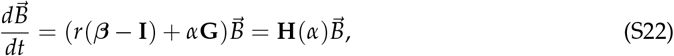

the last equality being the definition of **H**(*α*). On the assumption that the long-term behavior of the bacteria population is exponential growth or decay (as we have seen numerically), it follows that the steady-state cell distribution is the dominant eigenvector of **H**(*α*), and control is achieved when the dominant eigenvalue equals 0.

We observed numerically that the value of *α* required to achieve control is exactly *α* = *r*(1 + Λ). One way of proving this is based on the dynamics of total bacteria **B** = ∑_*i*_ *B*_*i*_,

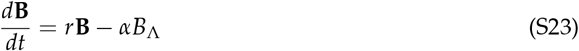

Zero population growth then occurs if and only if *x*_Λ_ ≡ *B*_Λ_/**B** = *r*/*α*. It is therefore sufficient to show that *x*_Λ_ = 1/(1 + Λ) when *α* = *r*(1 + Λ). Rewriting (S18) with *K* = ∞, and noting that 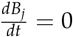 for all *j* implies 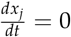 for all *j*, we obtain

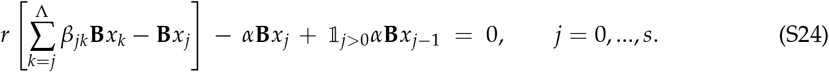

Dividing through by *α***B** and recalling that at equilibrium *x*_Λ_ = *r*/*α*,

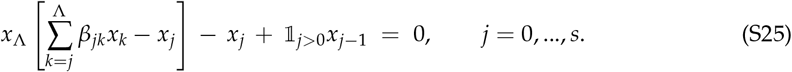

Multiplying each of the above equations by the corresponding *j* and adding them up,

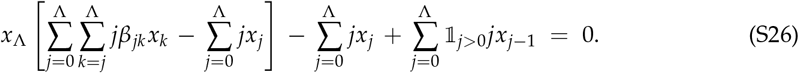

The content of the square brackets is equal to zero. To see this we can change the order of summation in the double sum:

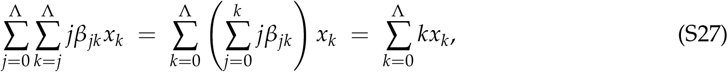

where the last equality follows from the standard property of binomial coefficients 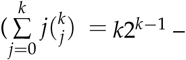 see also the “physical” explanation below). The remaining two terms in (S26) yield

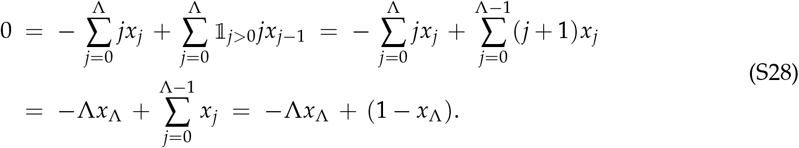

Thus, *x*_Λ_ = 1/(Λ + 1).

Another approach is to consider the growth of total “reproductive value”, 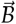 weighted by a left eigenvector of **H**. Let 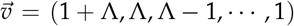 (a column vector) and **e** = (1, 1, …, 1) a column-vector equal in length to 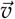. The following calculation was inspired by observing numerically that 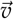 is a left eigenvector of **H**(*α*) when *α* = *r*(1 + Λ) with corresponding eigenvalue of 0.

We assert (and will prove below) that

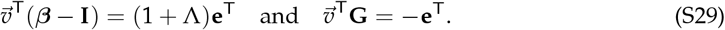

Accepting for the moment that (S29) is true, we have

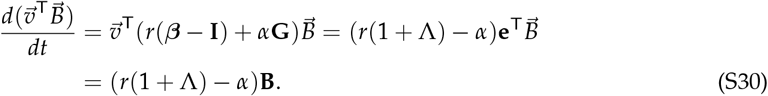

It follows from (S23) that 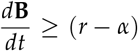 hence **B**(t) > 0 for all t whenever **B**(0) > 0. Therefore 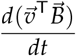 always has the same sign as r(1 + ^) − *a*. The proof is thus complete as soon as (S29) is proved.

The second identity in (S29) is easy. **G** looks like this:

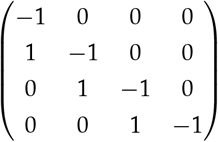

The first entry in 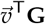 *v*^T^**G** is therefore −(1 + Λ) + Λ = −1; the second entry is −Λ + (Λ − 1) = −1; and so on until the last entry is last entry is -1, confirming that 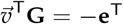.

The first identity in (S29) is not quite so easy. Start with the first entry, corresponding to zero hits. The first column of ***β*** is (2, 0, 0, … 0) because a parent with zero hits always produces 2 daughters with zero hits each when it divides. The first entry in 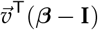 is therefore 2(1 + Λ) − (1 + Λ) = (1 + Λ), as claimed.

Other columns correspond to division of cells with a positive number of hits. Consider the column corresponding to a “parent” cell with *k* hits. The corresponding entry in 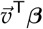 is

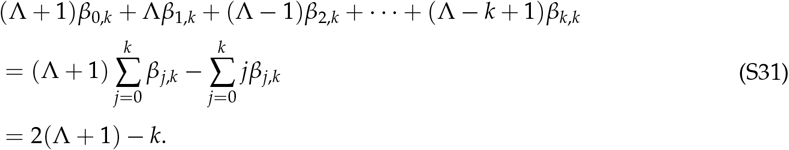

The last equality above follows from the “physical” interpretations of the sums. The first is the expected total number of daughters of a parent with *k* hits that divides, which is 2. The second is the expected total number of hits in all daughters of a “parent” cell with *k* hits, which equals *k*. And the final expression is equivalent to the first assertion in (S29).

Alternatively, the last equality above can be also derived without the “physical” interpretation, as follows. We have

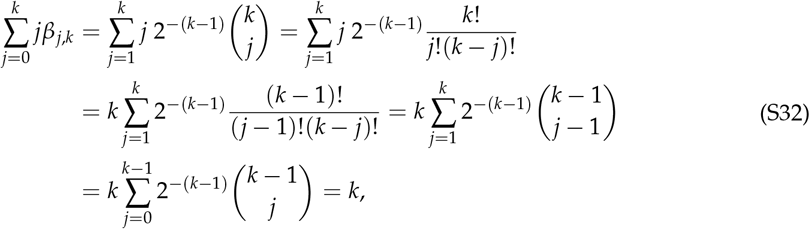

the last equality holding because the sum is the binomial expansion of (1/2 + 1/2)^*k*−1^ = 1.

### S7 Multi-objective optimization of host’s immune response

Suppose that *T* is a time horizon sufficiently long to determine the qualitative outcome of an infection (the infection is either controlled or the host is dead by the time *T* hours is reached). Assuming that the initial pathogen load and all model parameters are fixed, we consider two main metrics for the effectiveness and side effects of immune response.

- The total bacterial damage is assumed to be proportional to the time-integrated bacterial load,

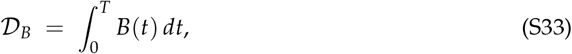

where *B*(*t*) is the total number of living bacteria at the time *t*.
- The total AMP auto-toxicity damage is assumed to be proportional to the time-integrated number of unattached AMPs,

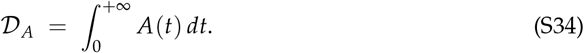

Both 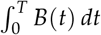 and 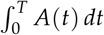 are easy to compute by augmenting the ODE system (1)-(9). However, the damage to host tissues from the AMP toxicity continues even after the infection is controlled, as long as there are any lingering AMPs. (This is why the upper limit in the second integral is +∞.) But based on our assumption that the host’s fate is determined long before the time *T*, it is reasonable to assume that no new AMPs are produced for *t > T*. If *B*(*T*) ≈ 0, the ODE for AMP numbers after that time are easy to solve analytically:

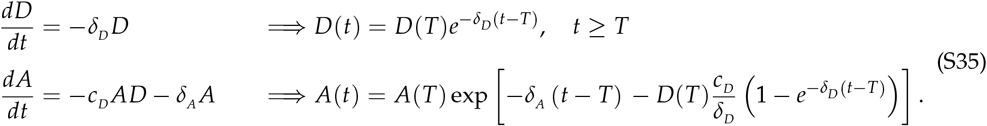

Using the above, the remaining AMP-toxicity damage 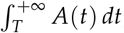, can be accurately approximated with the standard Gauss-Laguerre quadrature.

With bacterial-parameters and the initial load fixed, we can now explore how the immune response parameters could be adjusted (by evolutionary pressure) to minimize both the bacterial damage and the side effects. However, minimizing both 𝒟_*B*_ and 𝒟_*A*_ simultaneously is clearly impossible. Indeed, taking *V*_*A*_ = 0 ensures no toxicity damage (𝒟_*A*_ = 0), but will lead to a huge 𝒟_*B*_ and host death. Similarly, sending *V*_*A*_ → +∞ will control the infection with less and less bacterial damage, but will result in a dramatic increase in 𝒟_*A*_. Because these two optimization criteria are not aligned, it is necessary to consider all *Pareto optimal* regimes; i.e., the set of immune-response parameter values such that the resulting 𝒟_*B*_ cannot be decreased without increasing 𝒟_*A*_ (and vice versa). We accomplish this by using the standard *linear scalarization* approach. If **y** is a vector of optimized immune response parameters and *ξ* is an arbitrary number from [0, 1], we define

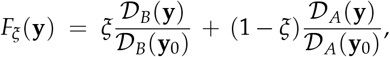

where both the bacterial and AMP damages resulting from **y** are normalized using the respective damages incurred with **y**_0_, the reference/default values of optimized immune response parameters. A standard argument (Miettinen, 1999) shows that, if 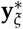 minimizes *F*_*ξ*_ (**y**) for any specific *ξ* ∈ (0, 1), then it must be Pareto optimal. (The coefficients *ξ* and (1 − *ξ*) can be interpreted as specifying the relative importance of these two types of damages.) This essentially reduces the problem to standard single-objective optimization: given a set of *ξ* values, we can minimize *F*_*ξ*_ for each of them and then display the set of 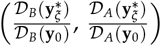 points, thus approximating (a portion of) the *Pareto front* for this bi-objective optimization problem; see Fig. 5 in the main text. The *ξ* values were first sampled from a regular coarse grid on [0, 1] and then more values were added adaptively as needed to accurately trace out the curve of Pareto optimal points.

Minimizing *F*_*ξ*_ (**y**) for a fixed *ξ* is a “standard” problem, but finding the global minimum can be computationally challenging when the function is non-smooth or rapidly changing. For example, near the “split” in outcomes (between controlled infection and host death), the gradient of *D*_*B*_ becomes extremely large, even though the optimal 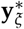 parameter values are nearby. This makes it harder to use methods that rely on smoothness of the objective function. In the text we consider two cases: **y** = *V*_*A*_ and **y** = (*V*_*A*_, *V*_*H*_). In the former, the one-dimensional minimization was performed using a combination of golden section search and parabolic interpolation (optimize function in R). In the latter two-dimensional case, we used nested one-dimensional optimization. Specifically the newuoa algorithm (Bates et al., 2023) was used to optimize *V*_*H*_, with the golden section method used to optimize *V*_*A*_ for every tested *V*_*H*_ value.

### S8 Effects of adjusting the dead cell clearance rate *δ*_*D*_

As we noted above in *§*S3, our default value for the dead cell clearance rate (*δ*_*D*_) is appropriate for our focal model system, adult *Drosophila melanogaster*, but other biological systems, including other insect species, invest more in clearance mechanisms. In this section we repeat the central analyses from the main text, allowing for higher values of *δ*_*D*_ on the infection time scale, and allowing *δ*_*D*_ to be one of the evolvable host defense parameters on the evolutionary time scale.

To represent more active clearance of dead cells and other debris, we chose a 10-fold increase in clearance rate, *δ*_*D*_ = 0.5, giving a half-life of roughly 1.4 hours. We also increased the relative AMP sponginess of the dead cells (*x*_*D*_) so that with default values of the other parameters, the model output remains very close to the split between host death and pathogen clearance after an initial dose of around 3000 live bacteria, in accordance with the empirical observations of (Duneau et al., 2017), and crosses the split as model parameters are varied around their default values. By trial and error, we found that a 2.5-fold increase in AMP sponginess (*x*_*D*_ = 2.5), which is still conservative relative to the experimental evidence (Savini et al., 2020; Snoussi et al., 2018)), compensated for the 10-fold decrease in the mean persistence time of dead cells.

Fig. S9 corresponds to Fig. 3 in the main text. The only qualitative change in behavior is that the fly dies within the simulated time period, rather than soon after, in scenarios where the immune response does not contain the infection and the bacteria population eventually exceeds the BLUD (Ω). This death of the fly results in the abrupt shutdown of AMP production. Apart from a slightly speedier demise of the doomed, the model behavior is qualitatively unchanged from the scenario in main text Fig. 3. Increasing the time-lag for immune up-regulation *τ*_*U*_ (panels A and B) again delays AMP production, allowing bacteria population growth to continue until the bacteria are so numerous that host’s AMP production cannot kill enough of them to prevent further increase. Higher sponginess of the dead (panels C-F) again has no detectable effect on

**Figure S9.**
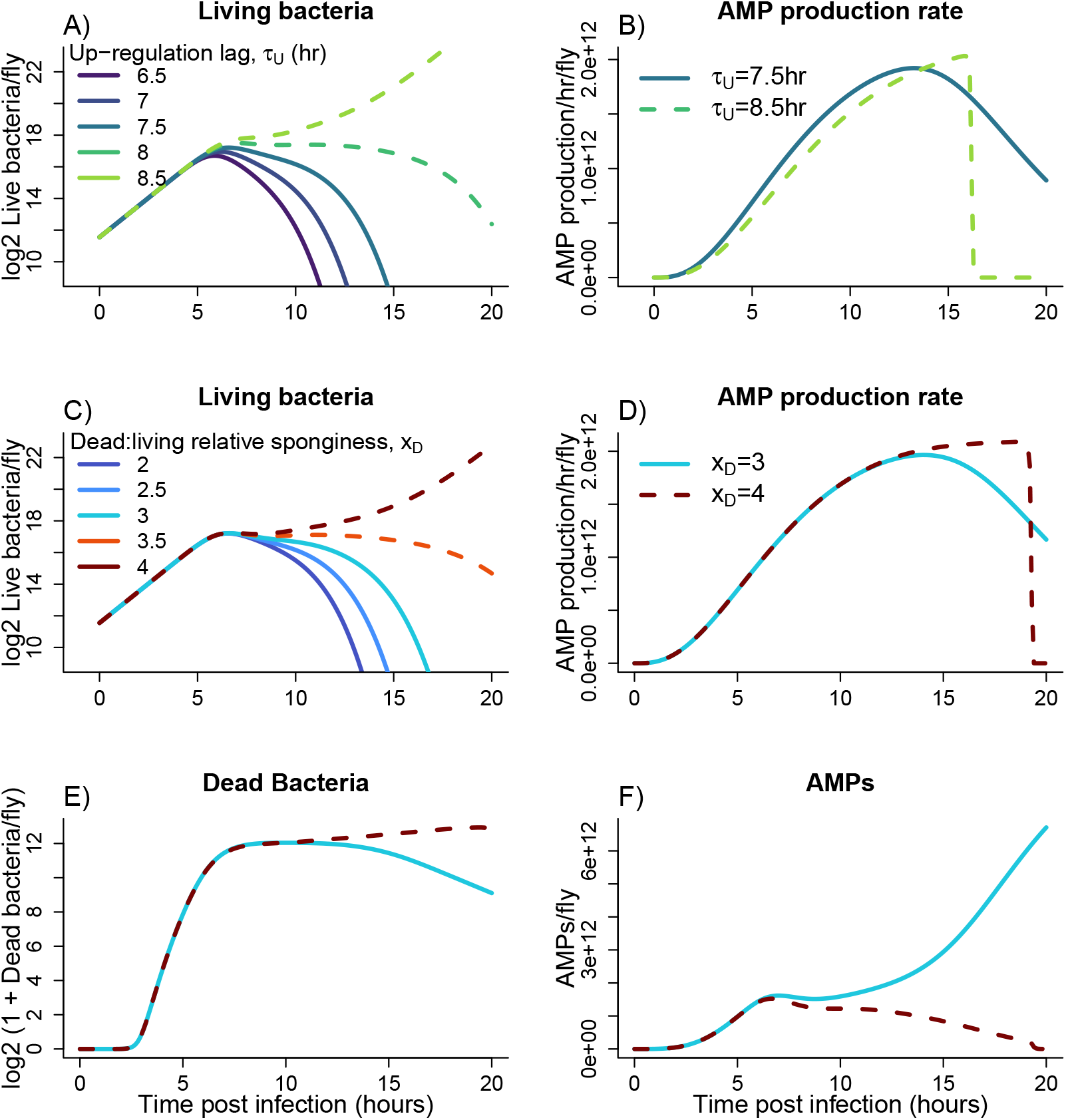
Re-creation of Figure 3 from the main text, assuming dead bacteria decay 10 *×* faster (*δ*_*D*_ = 0.5) and are more spongy (*x*_*D*_ = 2.5) relative to the default parameter values. Figure made by script `Figure3_moreSpongy.R`

AMP production, but once the dead are numerous, they sponge up so many AMPs that the living bacteria can continue to proliferate and kill the host.

Fig. S10 corresponds to Fig. 4 in the main text. Sensitivities are defined by small changes in individual parameters, close to the default parameter vector, so changing two of the default parameters (*δ*_*D*_ and *x*_*D*_) changes the sensitivities numerically. However, there are no changes in sign — no changes in whether increasing a parameter increases or decreases any particular cost.

**Figure S10.**
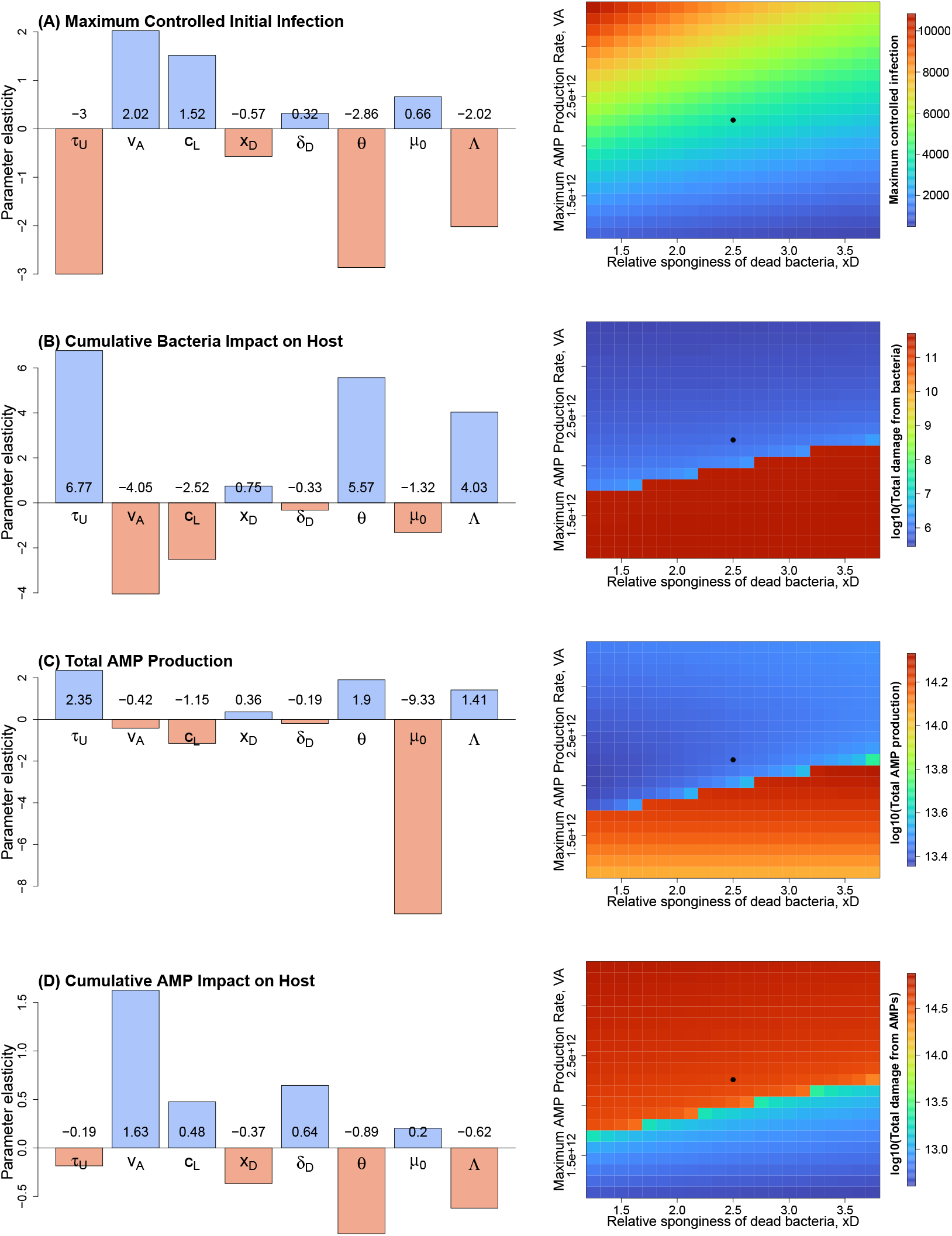
Re-creation of Figure 4 from the main text, assuming dead bacteria decay 10*×* faster (*δ*_*D*_ = 0.5) and are more spongy (*x*_*D*_ = 2.5) relative to the default parameter values. The main change in results is that because of increased dead sponginess, the response variables are more sensitive to parameters in the mortality function (*θ, µ*_0_, Λ). Panels made by scripts in the code archive Figure4_moreSpongy folder.

The main quantitative difference is that parameters in the bacterial mortality function (*µ*_0_ and *θ*) become substantially more important for most of the costs to the host. This change is intuitive; a larger value of *x*_*D*_ means that each bacterial death has a larger effect on the overall rate of AMP absorption by bacteria (living and dead combined) and therefore has a larger effect on the struggle between host and pathogen.

Because removal of AMP-sponging dead cells could contribute to a successful host defense, we also repeated our evolutionary time-scale analysis with *δ*_*D*_ rather than *V*_*H*_ as the second evolving parameter in tandem with the maximum AMP production rate, *V*_*A*_ . The (*V*_*A*_, *δ*_*D*_) pair includes one of the main parameters affecting the ramp-up of host defenses alongside another parameter affecting how long the bacterial corpses remain present and able to sponge the AMPs.

The main qualitative features in Fig. S11 are the same as those in Figure 5. Optimizing two parameters is still better for the host than optimizing one (i.e., the Pareto fronts move to the “southwest”, where both costs are lower, when two parameters are allowed to evolve). It also remains true that larger values of dead relative sponginess (*x*_*D*_) allow better outcomes for the host as the end result of defense parameter evolution. In line with the latter trend, if more AMP sponging by the dead benefits the host, the host also benefits from having the dead persist as long as possible — when *δ*_*D*_ is allowed to evolve, the optimized value of *δ*_*D*_ at all calculated points on the Pareto front is always *δ*_*D*_ = 0. That is: dead bacteria that persist forever will be maximally effective as the mop-up squad after the host attacks the pathogen with very rapid AMP production.

**Figure S11.**
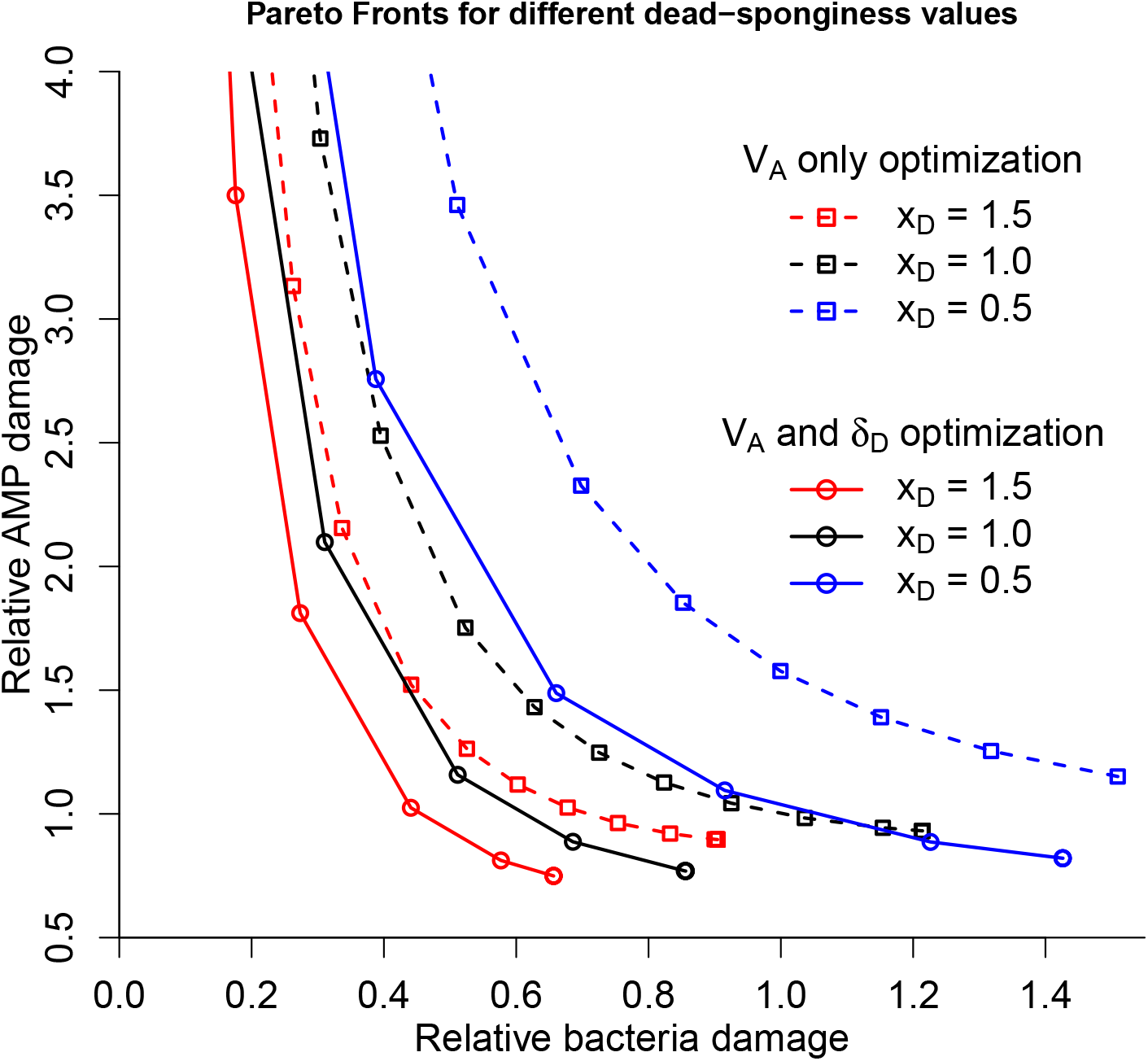
Re-creation of Figure 5 of the main text, in which the two evolvable parameters are *V*_*A*_ (maximum AMP production rate) and *δ*_*D*_ (the decay rate of dead bacteria). Dashed curves correspond to optimizing *V*_*A*_ only; solid curves correspond to optimizing both *V*_*A*_ and *δ*_*D*_ . Colors corresponds to dead bacteria “sponginess” relative to living bacteria (*x*_*D*_). The *V*_*A*_-only curves are drawn with points from Fig. 5 in the main text. The *V*_*A*_ and *δ*_*D*_ curves were generated by R script `Sponge_Optim_Adam_Bdam_Pareto_VA_deltaD.R` and the plot was made by Figure5_deltaD.R.

But the finding that in this computational experiment *δ*_*D*_ evolves to zero (meaning that the host should make no investment in mechanisms for clearing the dead) should not be regarded as a serious prediction about host immune evolution. Indeed, there are many insects that do make such investments (e.g., by having more circulating blood cells than adult *Drosophila* do). Instead, this observation re-emphasizes the point we make in the *Discussion* that taking account of more costs of defense for the host, such as the cost of AMP production, should be a priority for future investigations on the evolutionary time scale. Unfortunately, this will result in higher-dimensional Pareto fronts, which will be harder to visualize and use to evaluate the possible tradeoffs rigorously.

A deeper exploration of the effect on evolutionary time scales of mechanisms for clearing dead cells will also benefit from relaxing several of our simplifying assumptions:

1. Our model assumes immediate and complete release of PGN upon cell death, but in reality bacterial corpses probably release more PGN over time, having an ongoing immunostimulatory effect.
2. Even very high abundance of bacterial corpses is assumed to have no direct deleterious consequences for the host.
3. The host strategy of producing extremely high AMP concentrations and counting on dead bacteria to limit the autotoxic damage is only possible due to our simplifying assumptions that there is no intrinsic upper bound to the AMP production rate *V*_*A*_, and that maintaining the biochemical capacity to very rapidly produce AMPs is cost-free.
4. We have considered host defense against a single pathogen that the host can safely assume to be “spongy”. Relying on dead bacteria for clean-up after high AMP production may be too risky a strategy, if the dead cells of some pathogens have low sponginess. It may also be too risky if there is a chance of false positives (an immune response in the absence of an actual pathogen attack).

